# Explainable 3D CNNs Yield Progression-Linked Profiles of Saliency in Early Parkinson’s Disease Using T1-Weighted MRI

**DOI:** 10.1101/2025.11.17.688918

**Authors:** Ezekiel Moroze, Basilis Zikopoulos, Arash Yazdanbakhsh

**Affiliations:** Computational Neuroscience and Vision Laboratory, Department of Psychological and Brain Sciences, Boston University, Boston, MA, 02215 United States; Graduate Program for Neuroscience, Boston University, Boston, MA, United States; Center for Systems Neuroscience, Boston University, Boston, MA, United States; Human Systems Neuroscience Laboratory, Department of Health Sciences, Program in Human Physiology, Boston University, Boston, MA, 02215 United States; Department of Anatomy & Neurobiology, Boston University School of Medicine, Boston, MA, United States

## Abstract

Parkinson’s Disease (PD) affects 1-2% of adults over 65 and is diagnosed by tracking motor symptoms, underscoring the need for earlier detection. T1-weighted MRI (T1w MRI) is inexpensive and widely available but can lack the resolution to detect established markers of PD pathology, such as substantia nigra degeneration. We trained 3D convolutional neural networks (CNNs) on five site-stratified folds of 249 T1w MRIs from 141 early PD subjects (<10 years since first symptoms) and 185 from 126 controls. Because every fold was evaluated only on sites absent from its training data, the ensemble accuracy of 67.8% (AUC 0.670) reflects generalization to unseen sites. To interrogate these decisions, we parcellated saliency from a cohort-contrastive GradCAM, which baselines PD-class saliency against controls, and from Integrated Gradients, using the Julich Brain atlas. Both showed saliency in neocortical areas matching later Braak stages (5-6), when pathology reaches the neocortex, and lower subcortical saliency, suggesting limited sensitivity to earlier stages. Saliency was more correlated within functional networks than among random region groupings, and pooled saliency tracked motor symptoms and medication rather than cognitive scores or duration. These findings show explainable 3D CNNs can identify meaningful network and regional signatures of early PD from T1w MRI.

## Introduction

Parkinson’s Disease (PD) is a neurodegenerative disorder which affects between 1-2% of the global population over the age of 65 ^1,2^. Dysfunction and eventual death of dopaminergic neurons in the substantia nigra pars compacta (SNc) and associated areas is thought to be a primary neuropathological feature of early PD ^3,4^. This degeneration drives the motor symptoms characteristic of PD: bradykinesia (slowness of movement), tremor, and rigidity ^4,5^, and is further associated with decreased cognitive performance, changes in mood, decision-making, and sleep patterns ^5–8^. SNc atrophy cannot be reliably measured from widely available T1-weighted (T1w) MRI, which limits its value as a diagnostic tool ^9,10^. Instead, PD diagnosis centers around patient history and motor symptoms but may be supported by more specific imaging modalities like dopamine transporter (DAT) SPECT ^4,11,12^.

This limitation does not mean T1w MRI is uninformative about PD ^9^. Analyses of T1w MRIs have shown that cortical thickness changes in PD relative to healthy control in a way that correlates with cognitive decline^7,13,14^. Other T1w cortical thickness studies have observed a general spatial profile of cortical degeneration along PD progression, with thinning of frontal regions and the insula in the left hemisphere occurring in subjects less than 10 years from experiencing their first symptoms, and bilateral, widespread atrophy after 10 years ^15^. These findings indicate that T1w MRIs contain features of PD pathology across the disease’s time course, despite their difficulty resolving the SNc. That time course can be described by Braak staging, in which PD neuropathology spreads in a stereotyped histopathological sequence from the brainstem through subcortical structures to the neocortex, which gives a framework for asking how far pathology must advance before it leaves a signature detectable in T1w MRI ^16^. Given their worldwide availability, it would be advantageous to use big data approaches like deep learning to characterize how T1w MRIs reflect the effects of PD.

T1w MRIs have been used to train 2D convolutional neural networks (CNNs) to distinguish PD vs control (CT) at accuracies up to 70%, indicating there are discriminative features identifiable by deep learning models in structural imaging ^17^. 2D CNNs must be trained on MRI slices, meaning the models are unable to consider continuity across the entire MRI volume. 3D CNNs bypass this constraint by accounting for context across the entire brain in all three dimensions, granting an enhanced ability to recognize subtle spatial patterns and make accurate classifications when provided ample training data. While 3D models have the advantage in processing all features of MRIs, they require more training data than 2D because they cannot treat sections from a single MRI as multiple independent observations ^18^. Due to this limitation, sourcing enough images to train a 3D CNN for medical imaging tasks has been historically difficult ^18^. Fortunately, the creation of PPMI (Parkinson’s Progression Markers Initiative) has provided access to large, high-quality multi-center imaging datasets necessary to train effective 3D CNNs ^19–22^.

One such model is the 3D CNN proposed in Chakraborty et al. 2020. The 3D model outperformed 2D equivalents, reporting an average accuracy of 92.75% when classifying PD versus control on 406 MRIs from PPMI ^23^. While PPMI’s inclusion of multi-center data allows enough abundance for the training of CNN classifier models, sites remain heavily biased in the class of scan they contribute (PD, healthy control, etc.) ^20,22^. This means that failing to explicitly control for site between training and evaluation data can introduce class-correspondent signals rooted in site-specific noise. In fact, models perform worse when their evaluation is stratified on site: the model from Camacho et al. 2023 achieved an accuracy of 79.3% when isolating site in testing despite using a significantly larger dataset of 2,041 subjects. Reported accuracies are therefore sensitive to how strictly site is controlled, and an evaluation of how a model generalizes to unseen sites is likely to fall below estimates that do not isolate site.

As deep learning applications become more accurate and common in medical imaging analysis, it is critical to extensively analyze these complex models to unlock explanations for their classifications and avoid the “black box” pitfall commonly associated with AI ^24^. Making high-quality explainability a priority has been shown to increase trust among clinicians, a necessity if these tools ever enter clinical settings ^25^.

To address the issue of explainability in models, we employed a novel combination of explainability methods unlocked by our proposed 3D CNN, which discriminates early PD (<10 years since experiencing first symptoms) from control using 434 3T T1w MRIs from 267 subjects sourced from PPMI. We deployed a set of augmentations designed to combat multi-center variance and took advantage of transfer learning to help boost performance on our limited dataset. Evaluating every model on imaging sites it had never encountered in training, our five-fold ensemble reached an accuracy of 67.8%. By combining two gradient-based attribution methods, a cohort-contrastive form of GradCAM and Integrated Gradients, with a probabilistic anatomical atlas, we generated regional saliency maps indicating which parts of each MRI contributed most to its classification as PD or control, retaining only those regions whose saliency was reproducible across cross-validation folds. Correlation between saliencies of different regions implicated the nigrostriatal pathway and other canonical functional networks involved in PD, suggesting that structural data alone carries information aligned with functional organization. By relating saliency scores to subject motor scores, medication use, and disease duration, we connected our network’s explainable output to clinical data provided by PPMI and identified pooled regions and networks associated with symptoms and treatment in PD.

## Methods

### Dataset Curation and Preprocessing

Early Parkinson’s MRIs and subject data were sourced from the Parkinson’s Progression Markers Initiative (PPMI). We sampled T1-weighted MRIs with a 3.0T field strength from subjects between the ages of 55 and 75 who had experienced PD symptoms for under 10 years. The dataset was composed of 141 PD subjects and 126 control subjects from 39 different imaging sites. Some subjects contributed multiple MRI scans from different stages of their PD progression to the dataset; in total, the dataset comprised 249 PD scans and 185 control scans. Scans were from within the first 10 years since first experiencing symptoms. All MRIs were skull-stripped, intensity normalized and linearly registered to MNI152 space using FreeSurfer v7.3.2. Preprocessing quality was assessed via the Euler number reported by FreeSurfer, and images with a Euler number of -217 or below were excluded for significantly degraded image quality ^26^. To prevent cross-site leakage in our model evaluations, the dataset was split into a site holdout testing set as well as five site holdout validation sets. We extracted measures of motor symptoms (via Movement Disorders Society Unified Parkinson’s Disease Rating Scale Part 3, or UPDRS P3), cognitive scores (via Montreal Cognitive Assessment Scores, or MoCA), medication status (via Levodopa equivalent dose, or LEDD), and years since onset of symptoms for all scans and ensured they were balanced across folds as far as possible (Table 1). In the interest of including as much data as possible, we did not exclude subjects on the basis of comorbidities; comorbidities were nonetheless extracted and are reported in Supplemental Data 1, both for the entire dataset and for each per-fold validation set. See Figure 1 for a graphical overview of the study methods.

**Figure 1:**
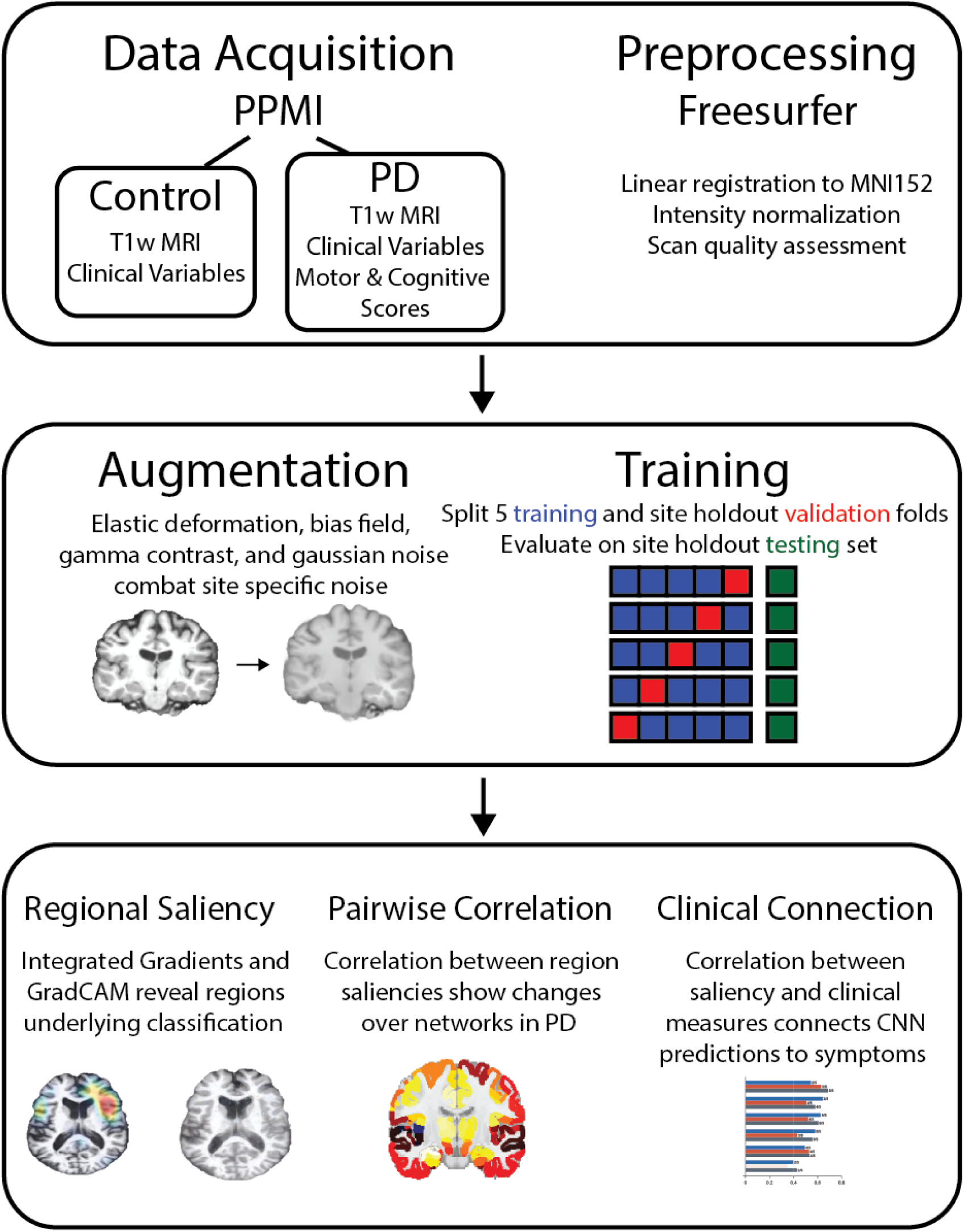
Graphical summary of study. Flow chart detailing how data was acquired and processed before training the model and how saliency data was interpreted in analysis.

**Table 1:**
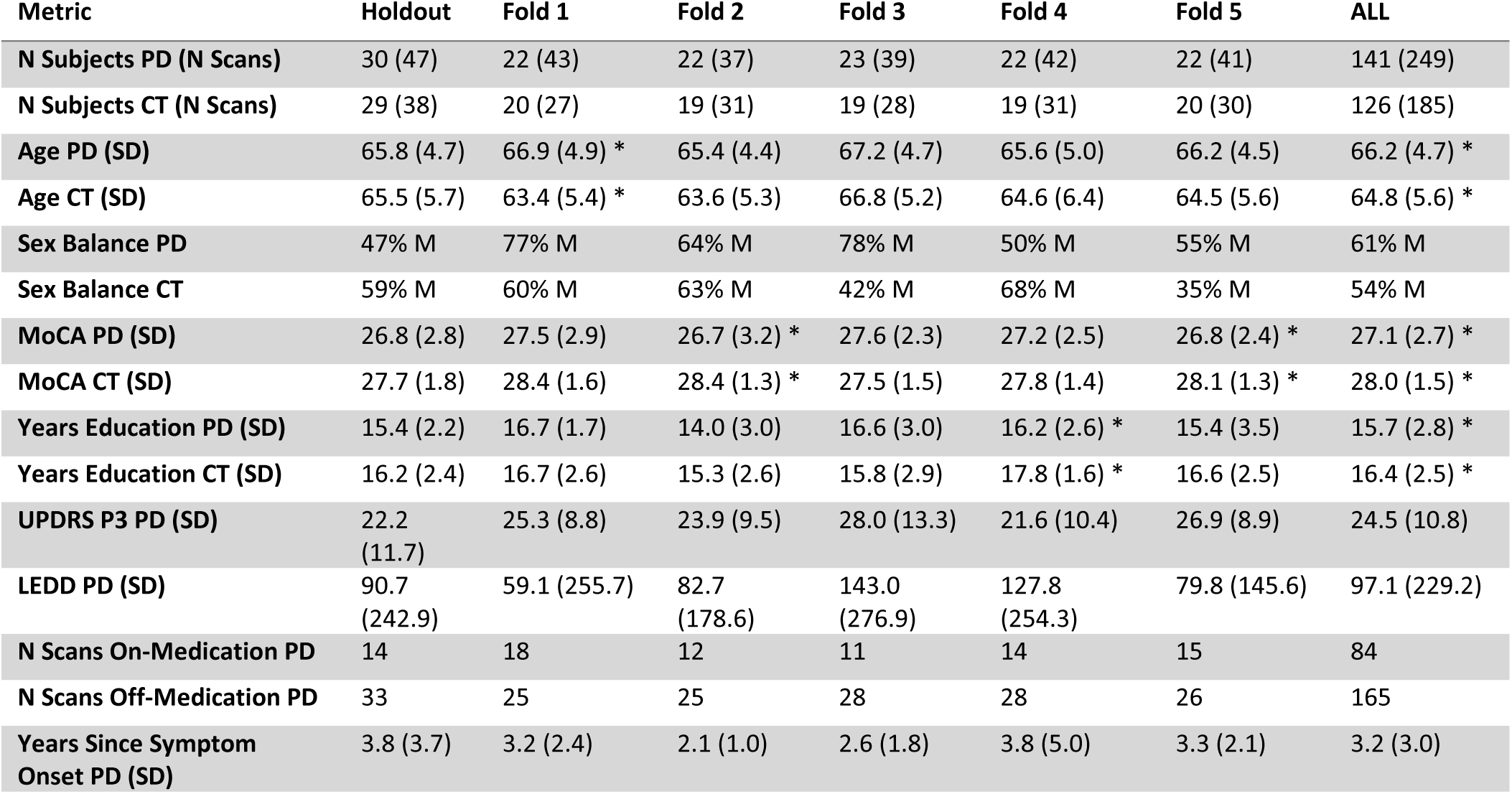
Breakdown of relevant subject metrics in dataset. Significant differences between PD and control (CT) are noted by asterisk (p < .05, Welch’s two tailed t-Test). We included multiple scans from some subjects. Accuracy, raw saliency, and pairwise saliency correlations are expressed at the subject level, while saliency-clinical correlations were computed at the scan level. Clinical attributes besides sex are reported at scan level, as they varied over multiple scans taken at different times for the same subject. The CT cohort was slightly younger than the PD cohort on average and had experienced less cognitive decline as measured by MoCA.

### Network Architecture

A ResNet-18 classifier pretrained on ImageNet (1.2 million images across 1,000 categories) was used as the base for our model. We selected a pretrained model to take advantage of a modified transfer learning paradigm, where our model was first pretrained in 2D on the large ImageNet dataset before being “transferred” to the PD vs CT task. In theory, this protocol serves to develop the ability to identify low level features (edges, basic shapes) using more abundant data to reduce the need to learn them on more limited datasets. In practice, this protocol has been shown to boost performance over unseeded networks in many classification domains, including medical imaging ^24,27^. To enable classification of MRI volumes instead of 2D sections while taking advantage of transfer learning, its weights were replicated along the z-dimension to convert 2D layers into 3D layers, then normalized by dividing by the size of the z-kernel ^28^. This means that, as initialized, the network would use pretrained 2D kernels to extract features along two slice planes of the MRI, and average local voxels in the remaining plane. Whereas early layers are traditionally frozen in transfer learning, we allowed them to train, but at a lower rate to ensure 3D relationships were learned across all anatomical planes while not overhauling transferred kernels. The learning rate progressively decreased with a scaling factor of 0.9 per residual layer going from deep to shallow (the shallowest residual block had a learning rate scalar of 0.48, compared with 1 for the deepest). To reduce overfitting by the deep layers, spatial dropout with probability 0.4 was added between convolutional units in residual branches. The pretrained fully connected layer was replaced with a 512×2 fully connected layer for binary classification of PD vs control. Biases were initialized to zero, and weights were initialized via Glorot initialization ^29^. See Figure 2 for a full schematic of the model architecture.

**Figure 2:**
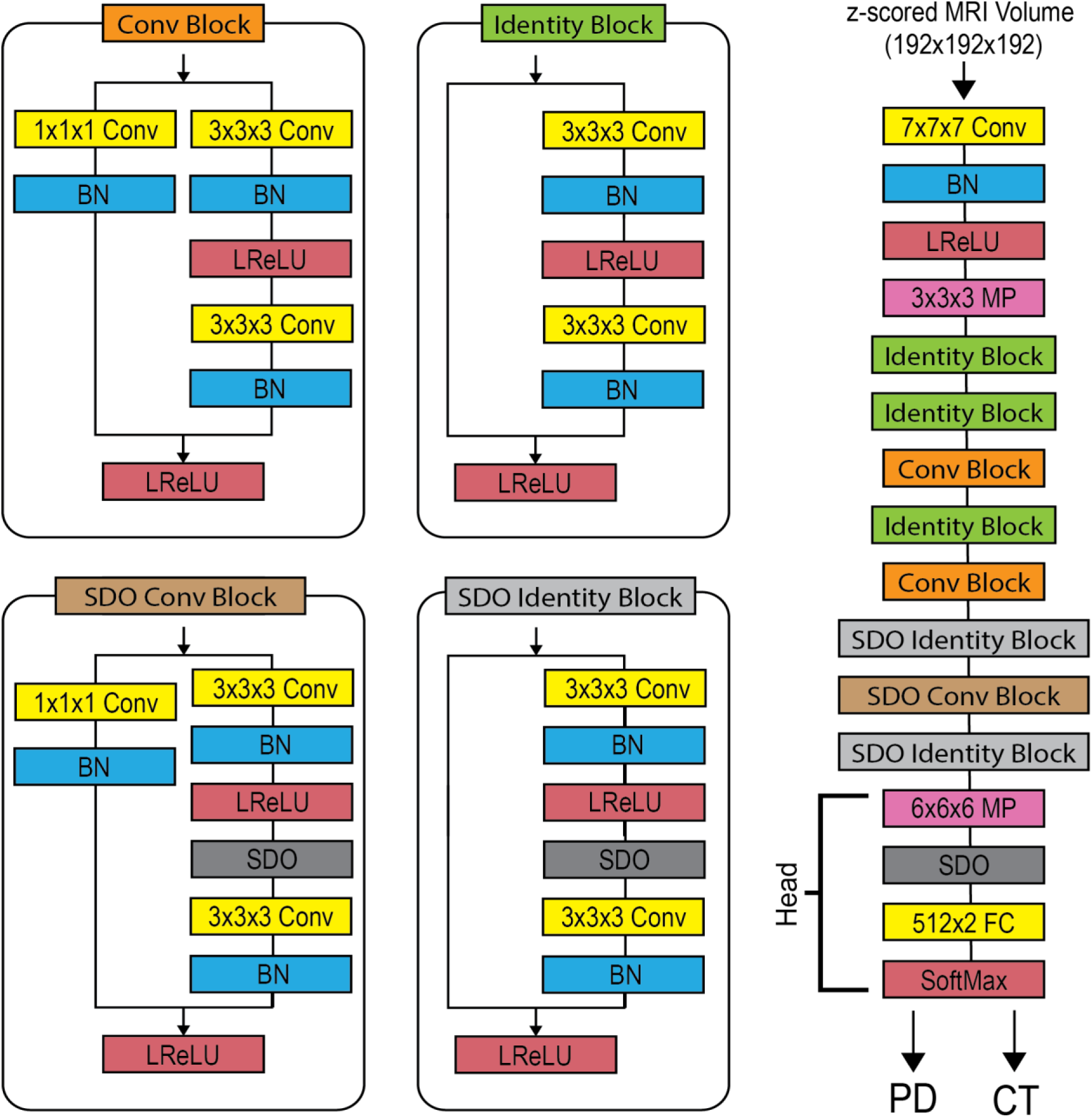
Modified 3D Resnet18 classifier architecture. The 3D CNN accepted z-scored 192x192x192 MRI images as input. Resnet is built from two different residual blocks, defined here as the identity block and conv block ^30^. Spatial dropout (SDO) was added to deeper residual branches. Classification was handled by a head segment containing a global average pooling layer (GAP), SDO with probability 0.4, fully connected layer (FC), and finally a SoftMax/classification layer which outputs final class scores. Other abbreviations: leaky rectified linear unit (LReLU); max pooling (MP), batch normalization (BN).

### Training & Reducing Site Specific Bias

PPMI is a dataset composed of images taken from a variety of imaging sites on a variety of scanner brands. This posed a problem, because CNNs have been shown to readily memorize subtle but systematic site-specific variance. Models may detect differences in bias fields or small geometric nonlinearities specific to a site’s particular scanner and acquisition pipeline ^31,32^. Furthermore, sites enrolled in PPMI did not contribute even numbers of PD and control subjects, with some even contributing subjects of a single class. Consequently, models can exploit site-specific signals as a shortcut, inferring pathology from site-level class imbalance rather than from anatomy. Thus, to obtain a more honest estimate of the models’ ability to detect generalizable anatomical signals in PD, we ensured that, for a given fold, sites in validation and test splits were always different from those in the training set.

The dataset had 267 total subjects; 141 PD and 126 control. We established a site-holdout test set of 59 subjects (30 PD, 29 control) with 85 scans (47 PD, 38 CT) to provide an unbiased estimation of classifier performance. Scans from the same subject were not always assigned the same class. Where they disagreed, the subject was given the label with the higher mean class score across that subject’s scans, and all performance metrics were reported at the subject level rather than the scan level to avoid individual subject bias based on the number of scans they contributed. The holdout set was composed of the 8 individual sites with the lowest class skew. We then divided the remaining 208 subjects into 5 folds to be used for 5-fold cross validation. The pretrained ResNet-18 described above was trained in MATLAB 2025a on each of the 5 folds, producing 5 fold-specific models. Each fold used the same hyperparameter set as determined by a grid search: ADAM optimizer, mini batch size of 10, 20 epochs, global learning rate of 3.0 * 10^-4^, shuffle data each epoch, and L2 regularization of 5.0 * 10^-4^. We took the weights from the lowest validation loss epoch for the final model for each fold. Following augmentation (described below), images were z-scored prior to entering the network to further regularize intensity histograms across sites.

The issue of site-specific variance was compounded by the fact that the 3D ResNet-18 had 33.2 million parameters but trained on only ∼166 subjects (∼280 scans) per fold. This imbalance meant the model would readily overfit, memorizing specific subjects rather than generalizable features of PD. While holding out sites for evaluation meant performance metrics were shielded from site specific leakage, it did not prevent the model from preferentially learning site-specific signals in the training set over subtler, generalizable PD features. To combat both issues, we deployed a comprehensive random augmentation set with the goal of masking (“harmonizing”) site specific variance and adding regularization to the limited training set. Augmentations were applied in the following order when a particular scan entered a minibatch for a training iteration: 50% chance of nonlinear elastic deformation (6x6x6 grid with displacement vectors randomly drawn from normal distribution of sigma 4 ^33^, linearly up sampled to input space and smoothed with spline filter), 90% chance of random bias field (scan multiplied by e to the voxel wise power of an upscaled 4x4x4 bias map sampled from normal with mean 1 and variance 0.25, then divided by the mean field strength), 70% chance of gamma contrast transformation with gamma uniformly drawn from a range of 0.61 to 1.65, 70% chance of additive Gaussian noise applied within the brain mask (variance 0.14 on a z-score scale).

### Saliency Methods, Atlasing & Masking

We used two different methods to extract regional explanations of classifier model decisions. GradCAM saliency maps provided coarse salience (or importance) maps upsampled to the whole subject MRI volume, in essence showing which regions drove each subject’s classification outcome (Fig 4, middle column). Integrated Gradients (IG) provided a map of network activations with voxel resolution by integrating the network gradient over a series of images interpolated between a control baseline and a PD subject’s MRI volume (Fig 4, right column).

**Figure 3:**
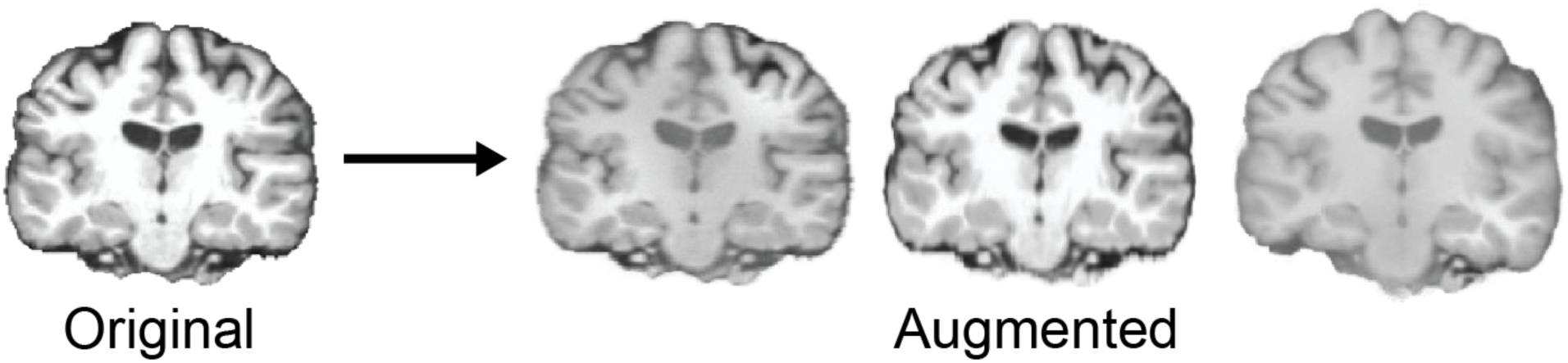
Examples of training volumes post random augmentation. All MRI volumes shown were sampled from the same PD subject. The original brain is how the brain appears after Freesurfer preprocessing. Three individual reads of the same subject are shown, showing the random changes in intensity, contrast, and subtle warping performed by the augmentation regime. The nondeterministic process helps prevent the CNN model from memorizing training data.

**Figure 4:**
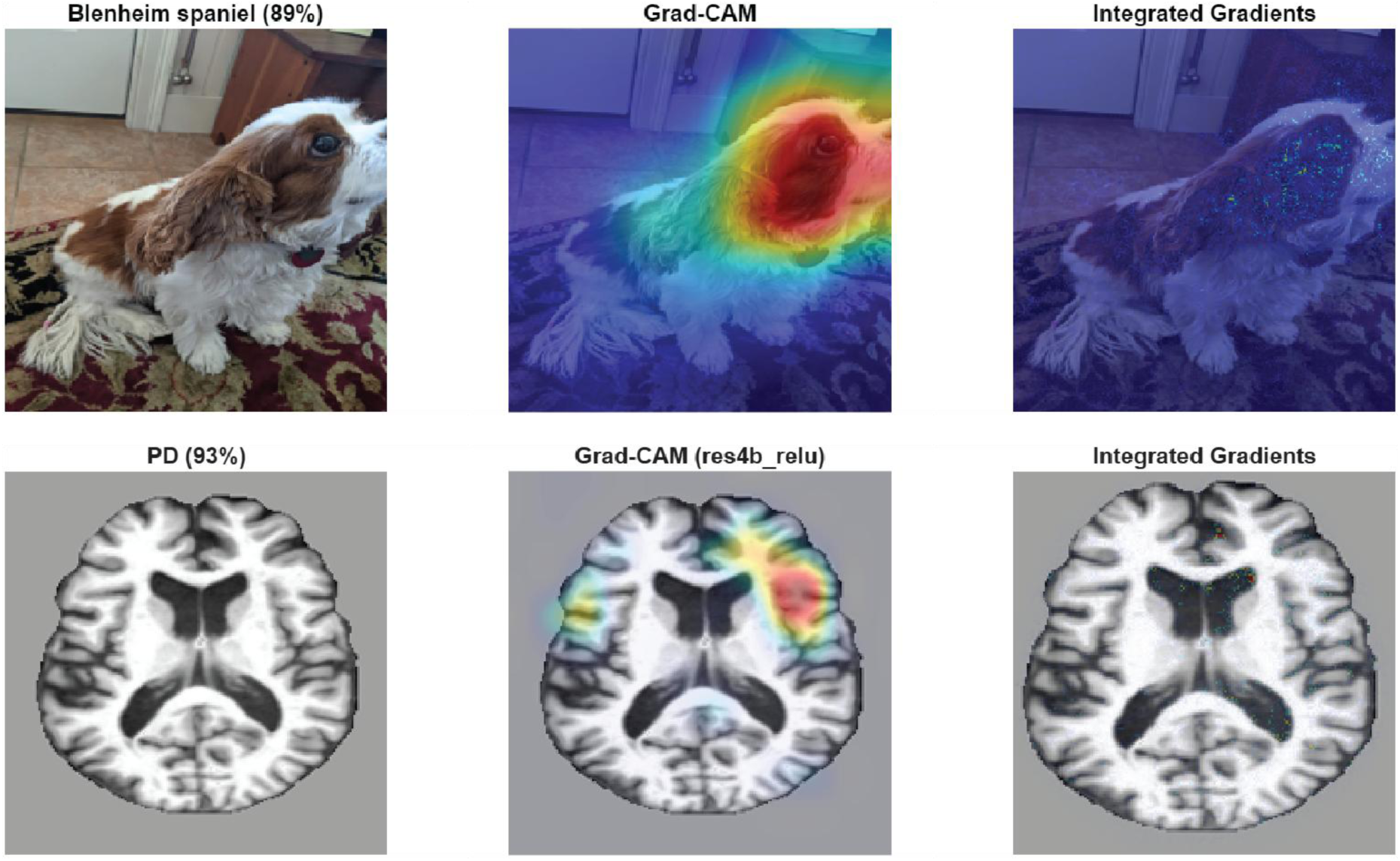
Examples of GradCAM saliency maps for different types of data. Top: example GradCAM and IG from class “Blenheim Spaniel” for an image of a dog, classified by a Resnet-18 trained on ImageNet. Percentage value next to class reflects model classification confidence. Res4b_relu refers to the CNN layer the GradCAM was sourced from. The GradCAM saliency map is more diffuse and shows the dog’s face and eye are salient features. IG tracks similar features but is less specific to the eye and spreads to the snout and ear with pixel resolution. Bottom: an axial slice from a PD subject’s GradCAM saliency map for class “PD”. Hotter regions for GradCAM can be interpreted similarly: the warmer parts of the brain are most informative to this subject’s PD classification. IG is much sparser and places subtle emphasis on the inside edge of ventricles and the frontal poles.

GradCAM works by taking the partial derivative of the target class logit for a given MRI with respect to a convolutional feature map, resulting in a channel weighting that indicates class-specific importance:

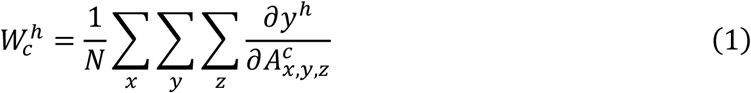

Where *W* is a weight corresponding to a specific channel *c* of the feature-map for binary class *h*. *W* is the sum of the partial derivatives of the class score *y^h^* with respect to a spatial feature 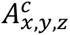 across all 3 dimensions. The final weight is normalized by *N*, the number of voxels in the feature-map. Finally, the channel-weighted sum of feature maps is half-rectified (ReLU) to give the gradient-weighted class activation map M which can be upsampled to the input space (known as the GradCAM saliency map) ^34,35^:

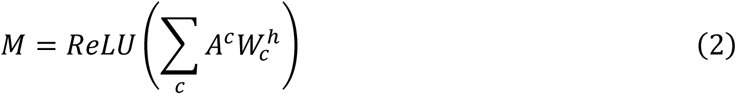

GradCAM enables the targeting of specific network layers for saliency map visualization. The deepest layers contain the most semantically relevant information, as they represent the features closest to where the final classification is made. However, they are also the lowest spatial resolution (our final two residual layers process features at a spatial resolution of 6x6x6). This makes the final two layers’ saliency maps too coarse relative to the fine scale of brain structures. We sampled all GradCAM maps from the third to last residual branch layer with a spatial resolution of 12x12x12, trading semantic specificity for spatial resolution.

This low resolution contrasts with Integrated Gradients, which computes voxel-wise attribution scores with respect to a target class logit along a linear path from a user-selected baseline to a specific input observation (a PD subject MRI) ^36^. The integral is approximated by the Riemann sum below:

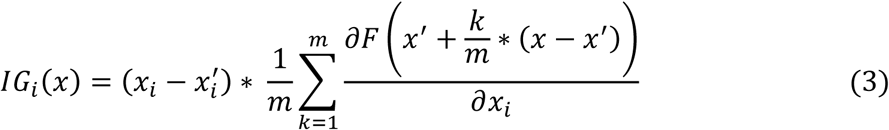

Where *x_i_* is a voxel of the image, 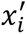 is the associated voxel in the baseline image, m is the number of steps in the Riemann approximation of the path integral, and F is the output of some CNN or other nonlinear function. We used *m* =200 steps to generate IG maps using the class logit from the fully connected layer.

IG satisfies the completeness axiom, meaning the sum of all voxel attributions equals the difference in logit between the input image and the baseline ^36^. Essentially, IG identifies how much a given voxel pushes the model toward or away from a target class relative to a baseline image. We selected this baseline image to ensure the IG signal would highlight features of PD that were discriminative relative to control. Taking inspiration from the Expected Gradients method ^37^, we calculated a given PD subject’s IG saliency map by averaging 25 IG maps, each using a random control subject unseen in training (testing set or that fold’s validation set) as baseline. This allowed for visualization of attributions from features present in the PD subject relative to the control brain while averaging out subject-to-subject anatomical variance unrelated to PD pathology.

Both IG and GradCAM saliency maps were generated from each scan per subject in the holdout testing set. Saliency scores were calculated for each of the 414 areas from the Julich Brain Probabilistic Atlas v3.1 by taking an element-wise, weighted average of each area’s probabilistic mask with a subject’s GradCAM and IG saliency volumes, computed independently for each method ^38^. This yielded, for each scan and each method, a vector of saliency values across the 414 atlas areas. For raw saliency and pairwise correlation analysis, we averaged together saliency scores for each region per subject. For more exploratory saliency clinical correlation, we used scan level regional saliency to correlate with clinical metrics derived from the date of the corresponding scan. Many metrics varied over time within subject, so this allowed us to boost the sample size of this analysis. The Siibra Python package was used to extract all probabilistic Julich atlas area names and masks in MNI152 space ^39^. A full list of atlas areas is available in Supplemental Data 2.

### Inter-fold Comparison and Validation

Because saliency data across subjects were generated from different network folds, it was critical to assess how similar saliency was across folds. As such, we calculated mean area saliency per area (across holdout test subjects) for each fold, then computed inter-areal Spearman correlations to assess how mean area-specific saliency scores aligned between folds, separately for IG and GradCAM (Fig 5).

**Figure 5:**
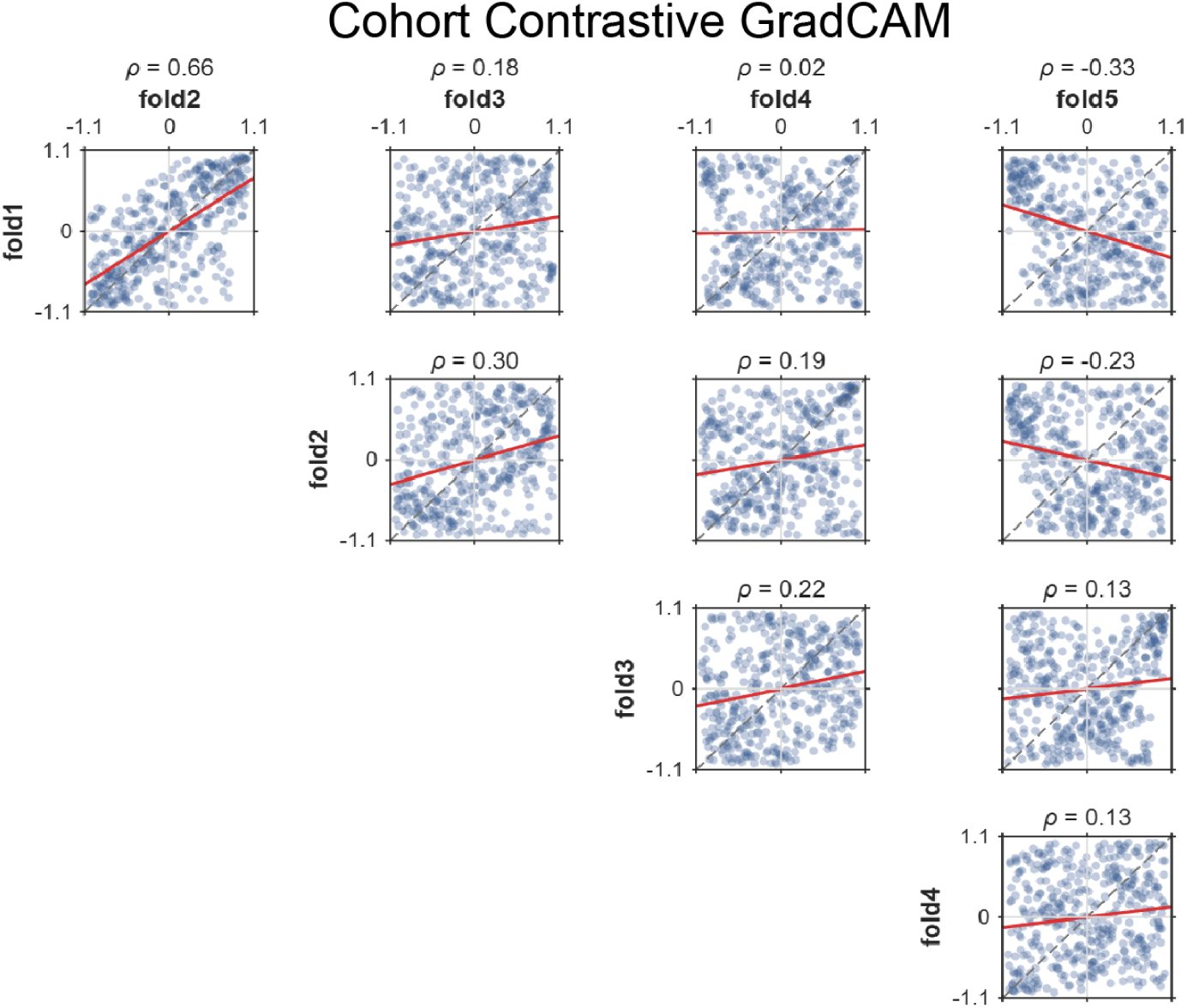
Cross-fold saliency concordance plots of average area saliency between all folds of the classifier model. GradCAMs were taken from the third to last residual layer, balancing resolution with semantic relevance. Each blue point per scatterplot represents an atlas region plotted by average saliency from different folds as aligned on the x and y axes. Alignment was measured via Spearman correlation. Due to the arbitrary units of saliency per network, each fold’s area saliency scores were independently rank normalized with sign preserved prior to plotting. Correlation between folds 1 and 4 was not significant (p < 0.05).

### Saliency Analysis

To establish what saliency scores might mean anatomically, we conducted a variety of analyses on the raw saliency scores. Firstly, we established a PD specific profile of raw saliency for both GradCAM and IG.

While IG was grounded to be discriminative relative to control via our multi-control subject baseline method, GradCAM lacked the same native class discriminative anchoring. This is complicated by the fact that GradCAM saliency has no concrete unit or common baseline, so GradCAM maps cannot be compared quantitatively across classes. However, GradCAM can calculate a saliency map for any target class, not just the predicted one, meaning we were in theory able to get a PD class GradCAM from a control subject. We used this capability to baseline the PD cohort’s GradCAM signal: for each region, we subtracted the PD-class GradCAM computed on control subjects from the PD-class GradCAM computed on PD subjects, leaving behind PD-discriminatory GradCAM signal. We used this cohort contrastive GradCAM signal (CCGCAM) to compare with control-baselined IG saliency.

To combine CCGCAM saliency across folds, we rank-normalized each fold’s CCGCAM saliency independently, as GradCAM has no universal scale or unit across different CNN models. The sign of CCGCAM was maintained through normalization to ensure we did not lose the class directionality CCGCAM affords. We filtered regions out by their robustness across folds to ensure we only reported reproducible signals. Because IG is signed, we defined robustness for IG as a given region’s saliency being the same sign across all five folds. For CCGCAM we applied a more stringent criterion: a region was robust only if saliency differed significantly between control and PD subjects (Benjamini-Hochberg adjusted Welch’s t-test, pFDR < .05) and if the difference was the same sign across all five folds.

While raw CCGCAM and IG saliency data provided a robust view of the regions which most consistently showed changes in PD, we wished to examine how saliency changes across pairs and networks of areas. Spearman correlations were computed across all correctly classified PD holdout subjects on a per-fold basis to circumvent GradCAM’s arbitrary unit scale. We filtered pairwise correlations by inter-fold robustness criteria: a pair had to be 1) the same direction in three of the five folds and 2) significant after FDR correction (pFDR < 0.05) in three of the five folds. If robust, the coefficients from all significant folds for that pair were averaged, and the mean coefficient was entered into a final inter-fold-robust 414×414 correlation matrix. We used this fold-robust correlation matrix to calculate the node degree for each region, which, for a given region *a*, was the sum of all correlation’s coefficients with all other regions *b* (Eq. 4). This metric provided a summary of how connected a region’s saliency was with that of others and treated both regions with many weaker connections and regions with fewer, stronger connections equally.

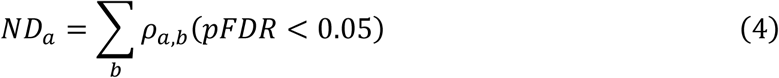

Where *a* is a given atlas area, *b* is the set of all atlas areas excluding *a*, *ρ_a_*_,*b*_ between a pair of areas represents significant (pFDR < 0.05) Spearman’s correlation coefficient. Because of the implicit strength-based weighting of the correlation coefficient, the node degree indicates an area saliency’s cumulative tendency to change with the saliency of others, meaning that high node degree areas may form central nodes or are linked with many areas in networks of saliency.

Given the possible impact of PD pathology on pathways/functional networks, we segmented our atlas into canonical functional networks described in literature (dorsal attention, ventral attention, executive/frontoparietal, visual, limbic, default mode, somatomotor) ^40–42^. Because they contain key PD ROIs such as the SNc, we also included nigrostriatal and mesocorticolimbic pathways in this analysis ^43,44^. We used these segmentations to compare the average correlation among areas within a functional network against the average correlation of those areas with areas in other networks. We used a region-label permutation test scheme to provide a more statistically rigorous comparison. In this method, for a functional network with N regions, we selected N random regions and measured their mean pairwise correlation. The permutation p-value was the fraction of random N-region sets whose within-vs-between difference was greater than the functional network’s difference after 10,000 trials. Supplemental Data 3 contains a full list of areas composing functional network segmentations.

Finally, we examined the correlation between coarse regional saliency measures and subjects’ scan clinical scores (UPDRS P3 total score for motor symptoms, MoCA for cognitive symptoms, LEDD for medication use, and the duration since first PD symptoms) for both GradCAM and IG. We extracted clinical variables from the time point of each scan, treating multiple scans within a subject as different observations. Because each of the metrics we were examining evolved over time within a subject, we could do this to boost the sample size of analysis. Instead of performing correlation on individual regions, we examined Pearson correlations over pooled regions defined by functional networks and lobes. This was done to reduce the number of correlations calculated and to control for the substantial intersubject noise in regional saliency. As a result, correlations were less fragile and more robust across folds, which was key to satisfying our reproducibility requirements. We used Pearson correlation for this analysis rather than the Spearman correlation applied elsewhere because pooling averages saliency across many areas, producing continuous and less skewed saliency distributions. Spearman correlation was preferred for the regional pairwise analysis because GradCAM saliency carries no meaningful unit, so only its rank ordering is comparable between networks. Despite performing this analysis at scan level, the sample size was still limited for poorly performing folds. As such, we relaxed our inclusion criteria and presented significant correlations in two or more folds.

## Results

### The Classifier Model

After training, the modified 3D ResNet-18 ensemble achieved an accuracy of 67.8% on our 59 subject (85 scan) testing set (0.708 F1, 0.67 ROC-AUC, Table 2). The ensemble model classified scans by determining if their average PD or CT score was higher across its five constituent folds. Where scans from the same subject were classified differently, the subject was assigned whichever label had the higher mean class score across their scans. We reported all accuracy at the subject level to avoid introducing bias from the number of scans a subject contributed (Supplemental Data 4 shows how many subjects had diverging scans per fold). This ensemble voting improved accuracy over any individual constituent fold (the next highest was fold 4, at 66.10%). Three folds (2, 3, 4) performed near chance on their validation sets, but this could be in part due to heavy site biases in their validation set composition (most of folds 3 and 4’s validation subjects came from a single site; Supplemental Data 5).

**Table 2:** Per fold and ensembled 3D Resnet18 performance on binary PD classification task. Validation set performance was used to determine the early stopping point for each network, but validation sets were not as well site-balanced and so were not analyzed further. See Supplemental Data 5 for statistics of site makeup per fold validation set. All metrics are reported at the subject level to avoid biasing metrics based on how many scans a subject contributed. Where scans from the same subject were classified differently, the subject was assigned the label with the higher mean class score across their scans. ROC-AUC: receiving operating characteristic-area under curve.

|  | Fold 1 | Fold 2 | Fold 3 | Fold 4 | Fold 5 | Ensemble |
|---|---|---|---|---|---|---|
| Validation Accuracy | 63.40% | 53.70% | 54.80% | 55.00% | 66.70% | -- |
| Test Accuracy | 64.40% | 54.20% | 64.40% | 66.10% | 57.60% | 67.80% |
| Test F1 | 0.656 | 0.571 | 0.553 | 0.737 | 0.49 | 0.708 |
| Test ROC-AUC | 0.686 | 0.553 | 0.638 | 0.587 | 0.685 | 0.67 |

Following model testing and evaluation, we examined if any clinical attributes or comorbidities distinguished the misclassified subjects from the properly classified subjects in the test set. The only significant difference between groups was the age between correctly and incorrectly classified controls, with correctly classified controls being on average nearly 6 years younger than incorrectly classified controls (62.85 years old vs 68.78 years old, pFDR < 0.001, Supplemental Data 6). This suggests the network conflated age-related change with PD pathology to some degree.

Folds varied considerably in their test set performance. For CCGCAM and IG we examined cross-fold saliency concordance to see if different folds were using different regions as basis for their classifications (Fig 5, 6). Overall, both CCGCAM and IG had limited interfold alignment, with mean off-diagonal Spearman coefficients of 0.121 and 0.126, respectively (Fig 5, 6 top). Overall, folds showed low concordance in regional saliency, indicating a need to stringently filter regions by fold agreement to reduce noise and to ensure signal was replicable. However, when fully rectified, IG’s Spearman correlation increased to 0.456 (Fig 6 bottom). Because a majority of IG’s signal clustered around zero, differences in saliency signs likely caused a high degree of volatility between folds, which full rectification removed. Given this observation, a region was filtered out from IG saliency visualization if any of the five folds disagreed on the sign of saliency. A region was filtered from CCGCAM if any of the five folds disagreed in sign of the difference between PD and control cohorts or if there was no significance.

**Figure 6:**
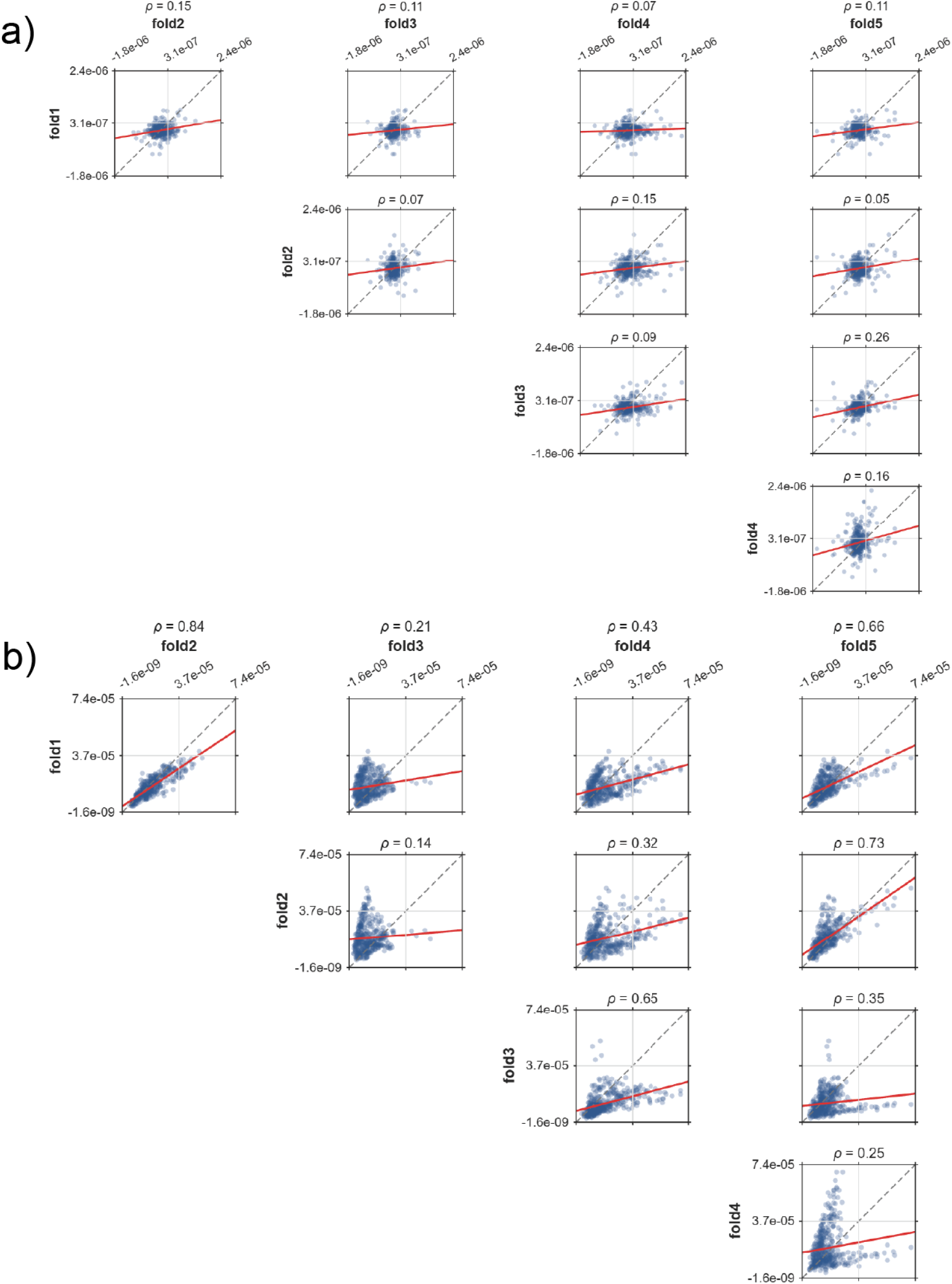
Cross-fold saliency concordance plots of average area saliency between all folds of the classifier model. Integrated Gradients (IG) maps were averaged across 25 different maps using 25 different control subjects as baseline images, giving an approximate class contrastive signal. Panel a shows concordance plotted according to mean saliency of the raw IG signal; correlations between folds 1 and 4, folds 2 and 3, folds 2 and 5, and folds 3 and 4 were not significant (p < 0.05) for mean IG. Panel b is plotted against mean of fully rectified (FR) IG signal. All pairs of correlations were significant for FR IG. Each blue point per scatterplot represents an atlas region plotted by average saliency from different folds as aligned on the x and y axes. Due to IG’s completeness, between fold saliency was not normalized.

### Raw Class Contrastive Saliency Scores

To understand the regional profile of saliency which distinguished the PD class from control, we visualized both CCGCAM signal (Fig 7) and IG signal (Fig 8) and examined which regions were robust across folds and attribution methods (Table 3). Several regions in the left frontal lobe showed fold and method robust saliency profiles, including parts of the pre-supplementary motor area (preSMA) and the frontal pole. Orbitofrontal and frontal opercular areas were robust but differed in sign between CCGCAM (CT>PD) and IG (towards PD). This divergence between methods could also be observed in other regions in and around the right operculum, such as the right parietal operculum and the right inferior parietal lobule. In the left hemisphere, regions around the operculum tended to be very high in saliency across both methods, suggesting these were the regions the model relied on most when making classifications. While nearly all robust subcortical areas were left lateralized, only the ventral striatum agreed in direction (toward PD, PD > CT) between our two saliency methods. Others (including the left SNc and thalamic parcellations) showed the same diverging pattern as in the right operculum where IG showed signal towards PD and CCGCAM showed greater saliency for control subjects than PD subjects for the PD class.

**Figure 7:**
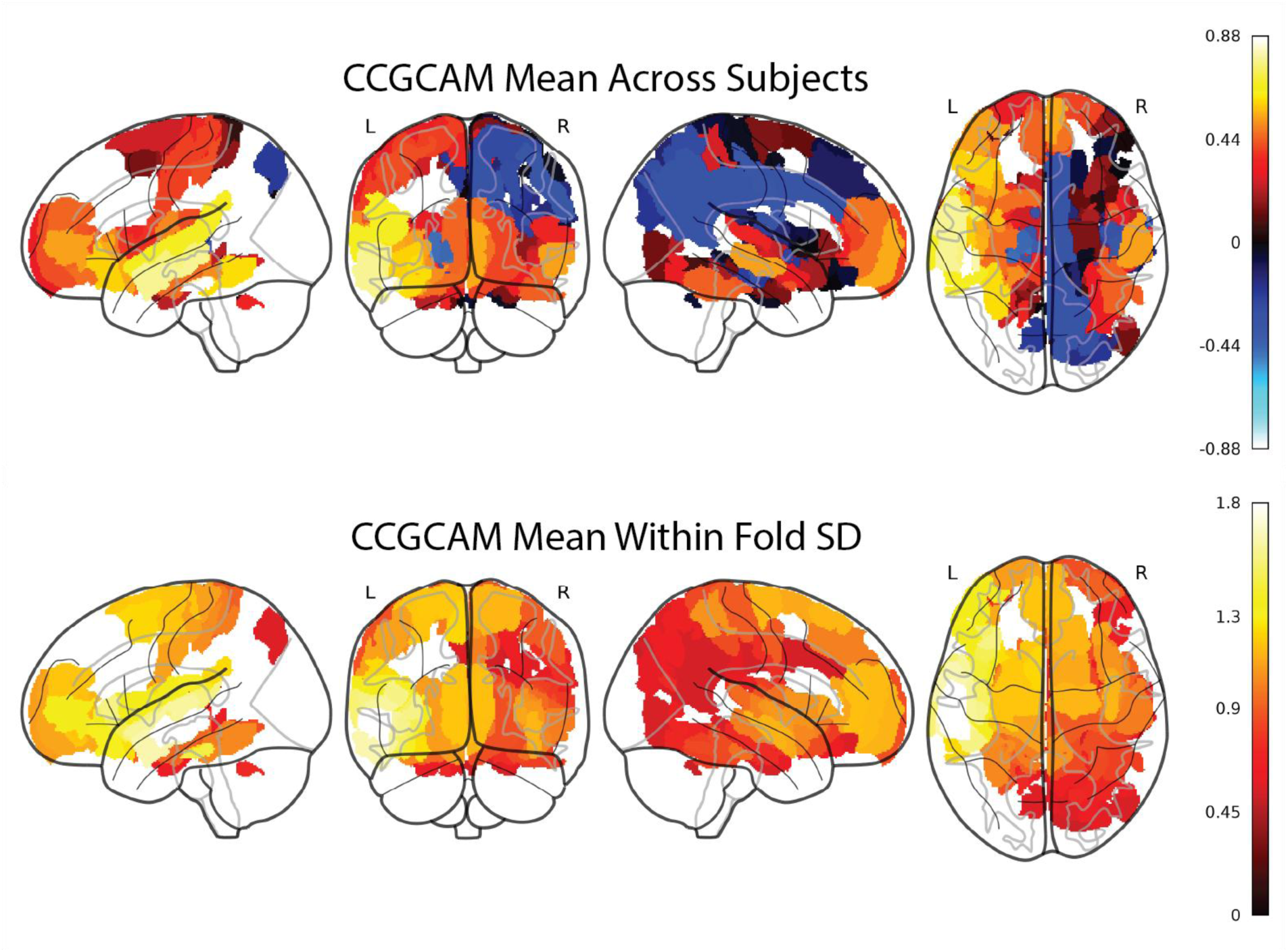
Cohort contrastive GradCAM (CCGCAM) saliency profile for PD subjects. Visualizations show brain regions colored by the difference in saliency for PD class GradCAM for PD subjects and PD class GradCAM for control subjects averaged across folds (cohort contrastive GradCAM or CCGCAM, top). Also visualized is regional within-fold standard deviation (SD) across subjects averaged across folds to visualize intersubject variability (bottom). CCGCAM signal was rank normalized with sign maintained per fold before averaging and SD calculation. Because high regional variance across folds was observed, regions were only included in visualization if 1) PD class PD subject and PD class control subject class saliency was significantly different across all five folds (Welch’s t-test, pFDR < 0.05) and 2) the difference between PD and control subjects was the same sign for that region across all five folds. Standard deviation (SD) across correctly classified subjects was pooled across folds and is represented on bottom. Left (L) and right (R) hemispheres are labeled over brain volumes.

**Figure 8:**
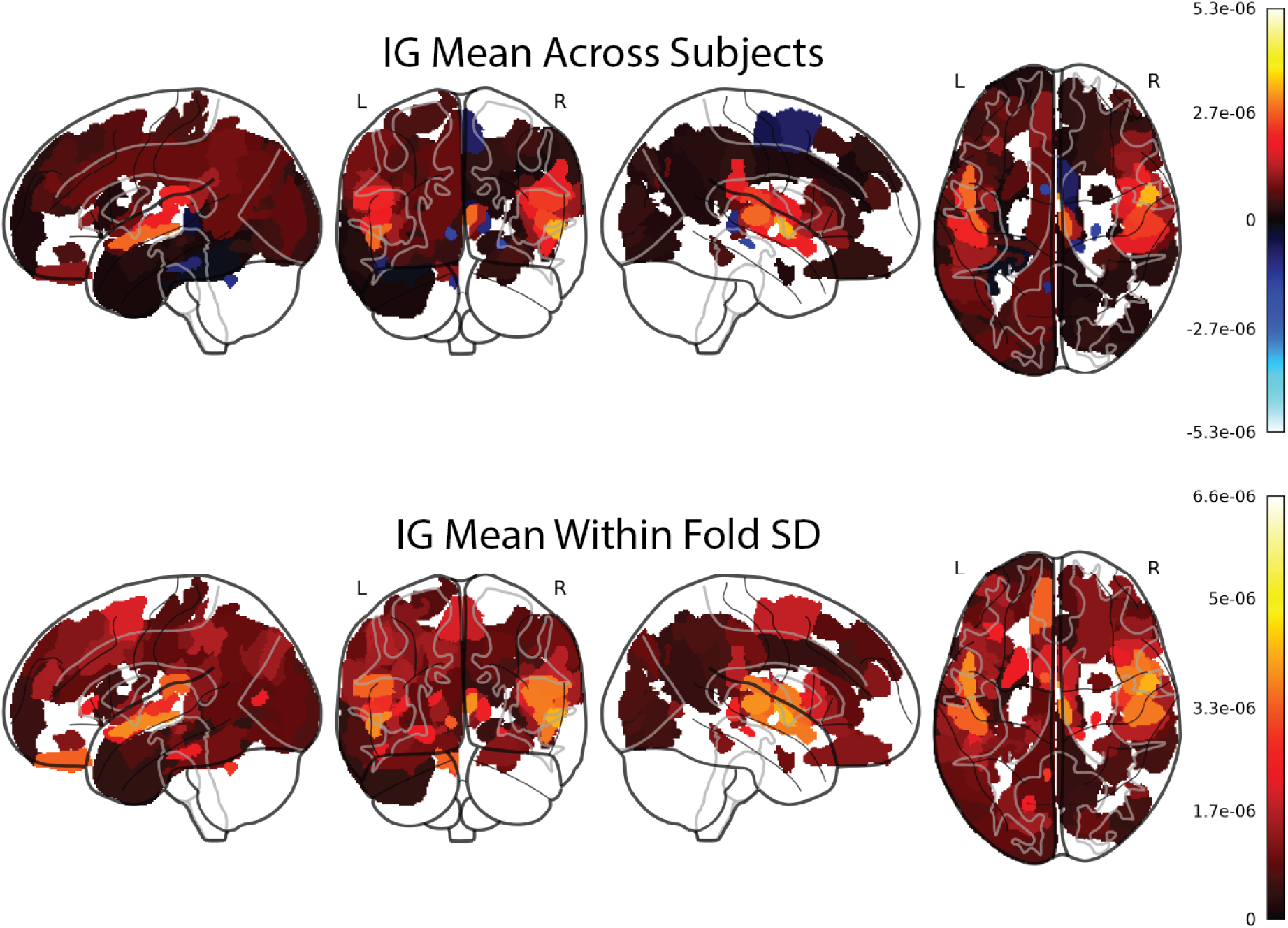
Integrated Gradients (IG) saliency profile for PD subjects. Visualizations show brain regions colored by the regional IG signal averaged over 25 random control subject baselines to control for noise due to intersubject anatomical variance not correspondent to PD pathology. Because of IG’s noisy, high volatility in saliency sign around zero, regions were only included in visualization if all five folds agreed in the sign of saliency. Standard deviation (SD) across correctly classified subjects was pooled across folds and is represented on bottom. Left (L) and right (R) hemispheres are labeled over brain volumes.

**Table 3:**
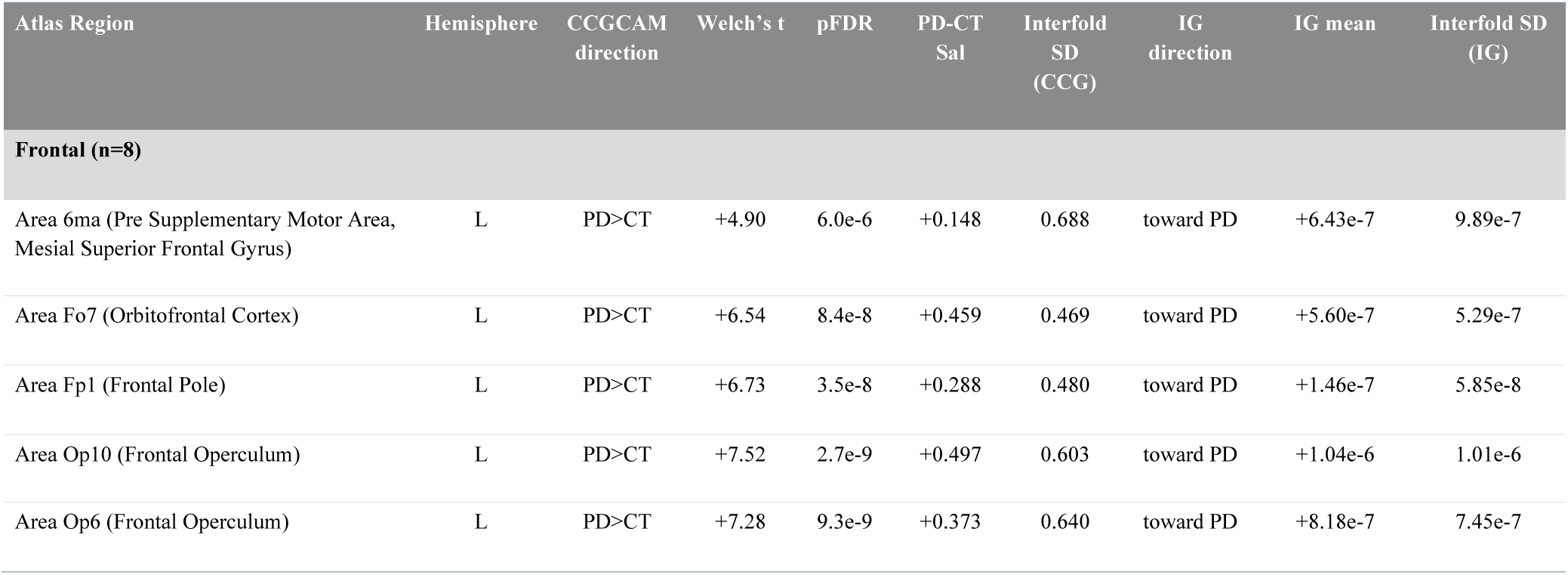
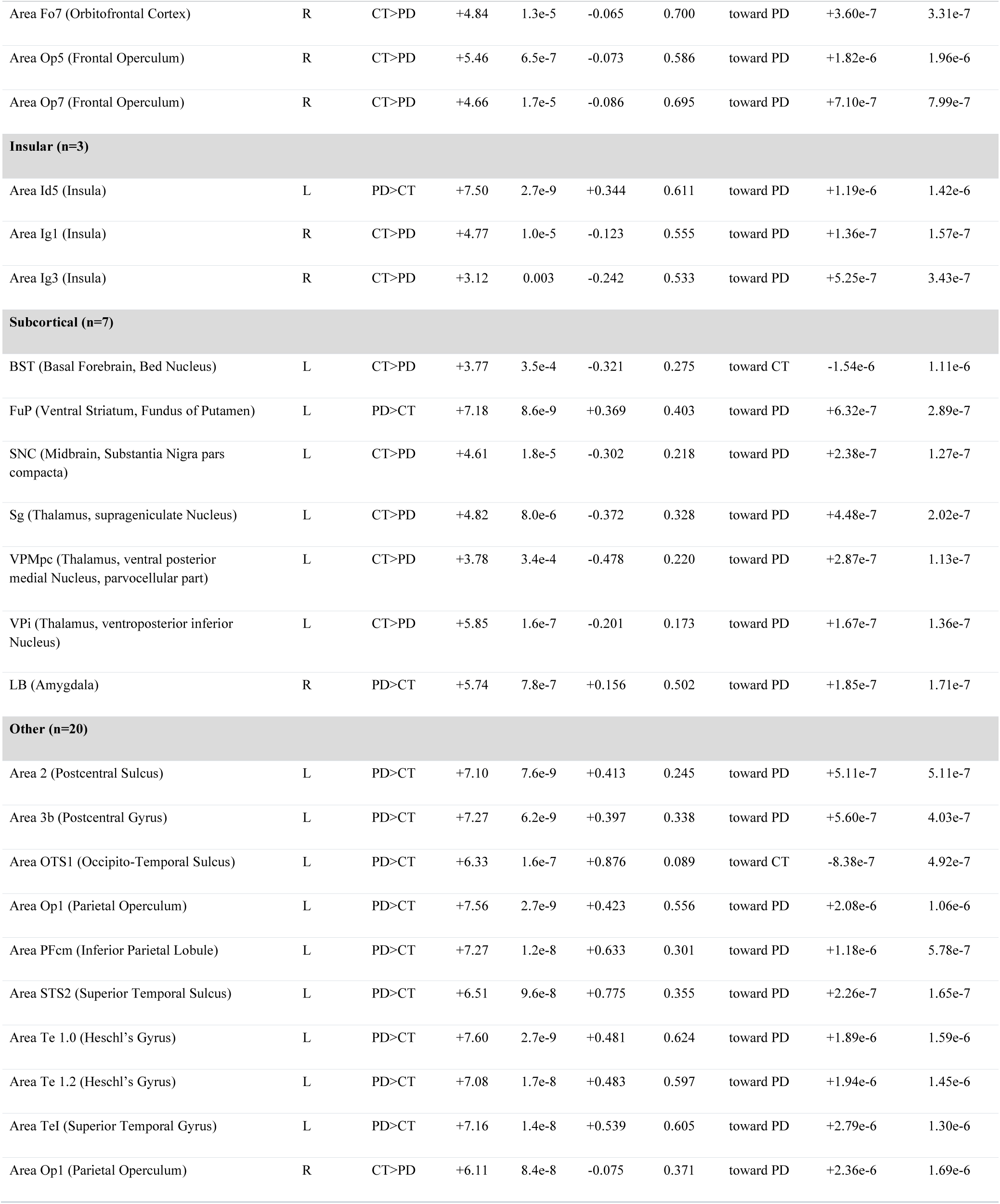
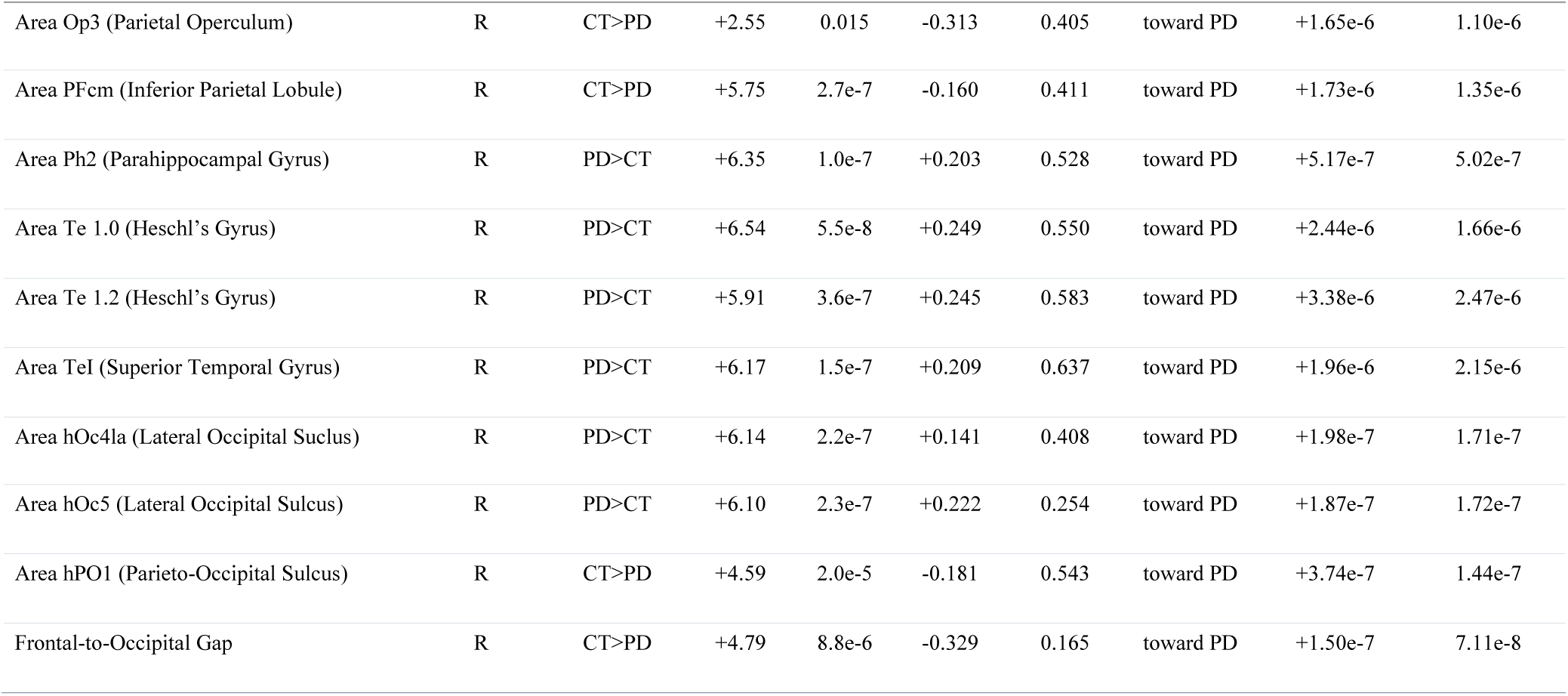
Regions with robust/reproducible saliency for both cohort contrastive GradCAM (CCGCAM) and Integrated Gradients (IG). All 38 atlas regions which survive both IG and CCGCAM interfold robustness criteria (sign match and difference significant across all five folds for CCGCAM, sign match across all five folds for IG) are presented with cohort direction (CCGCAM direction), pooled Welch’s t-statistic, BH-FDR adjusted pFDR value, CCGCAM saliency after per fold rank normalization (PD-CT Sal), standard deviation across all five folds. IG direction, IG mean saliency, and standard deviation across all five folds is also reported for comparison.

| Atlas Region | Hemisphere | CCGCAM direction | Welch’s t | pFDR | PD-CT Sal | Interfold SD (CCG) | IG direction | IG mean | Interfold SD (IG) |
|---|---|---|---|---|---|---|---|---|---|
| **Frontal (n=8)** |  |  |  |  |  |  |  |  |  |
| Area 6ma (Pre Supplementary Motor Area, Mesial Superior Frontal Gyrus) | L | PD>CT | +4.90 | 6.0e-6 | +0.148 | 0.688 | toward PD | +6.43e-7 | 9.89e-7 |
| Area Fo7 (Orbitofrontal Cortex) | L | PD>CT | +6.54 | 8.4e-8 | +0.459 | 0.469 | toward PD | +5.60e-7 | 5.29e-7 |
| Area Fp1 (Frontal Pole) | L | PD>CT | +6.73 | 3.5e-8 | +0.288 | 0.480 | toward PD | +1.46e-7 | 5.85e-8 |
| Area Op10 (Frontal Operculum) | L | PD>CT | +7.52 | 2.7e-9 | +0.497 | 0.603 | toward PD | +1.04e-6 | 1.01e-6 |
| Area Op6 (Frontal Operculum) | L | PD>CT | +7.28 | 9.3e-9 | +0.373 | 0.640 | toward PD | +8.18e-7 | 7.45e-7 |
| Area Fo7 (Orbitofrontal Cortex) | R | CT>PD | +4.84 | 1.3e-5 | -0.065 | 0.700 | toward PD | +3.60e-7 | 3.31e-7 |
| Area Op5 (Frontal Operculum) | R | CT>PD | +5.46 | 6.5e-7 | -0.073 | 0.586 | toward PD | +1.82e-6 | 1.96e-6 |
| Area Op7 (Frontal Operculum) | R | CT>PD | +4.66 | 1.7e-5 | -0.086 | 0.695 | toward PD | +7.10e-7 | 7.99e-7 |
| **Insular (n=3)** |  |  |  |  |  |  |  |  |  |
| Area Id5 (Insula) | L | PD>CT | +7.50 | 2.7e-9 | +0.344 | 0.611 | toward PD | +1.19e-6 | 1.42e-6 |
| Area Ig1 (Insula) | R | CT>PD | +4.77 | 1.0e-5 | -0.123 | 0.555 | toward PD | +1.36e-7 | 1.57e-7 |
| Area Ig3 (Insula) | R | CT>PD | +3.12 | 0.003 | -0.242 | 0.533 | toward PD | +5.25e-7 | 3.43e-7 |
| **Subcortical (n=7)** |  |  |  |  |  |  |  |  |  |
| BST (Basal Forebrain, Bed Nucleus) | L | CT>PD | +3.77 | 3.5e-4 | -0.321 | 0.275 | toward CT | -1.54e-6 | 1.11e-6 |
| FuP (Ventral Striatum, Fundus of Putamen) | L | PD>CT | +7.18 | 8.6e-9 | +0.369 | 0.403 | toward PD | +6.32e-7 | 2.89e-7 |
| SNC (Midbrain, Substantia Nigra pars compacta) | L | CT>PD | +4.61 | 1.8e-5 | -0.302 | 0.218 | toward PD | +2.38e-7 | 1.27e-7 |
| Sg (Thalamus, suprageniculate Nucleus) | L | CT>PD | +4.82 | 8.0e-6 | -0.372 | 0.328 | toward PD | +4.48e-7 | 2.02e-7 |
| VPMpc (Thalamus, ventral posterior medial Nucleus, parvocellular part) | L | CT>PD | +3.78 | 3.4e-4 | -0.478 | 0.220 | toward PD | +2.87e-7 | 1.13e-7 |
| VPi (Thalamus, ventroposterior inferior Nucleus) | L | CT>PD | +5.85 | 1.6e-7 | -0.201 | 0.173 | toward PD | +1.67e-7 | 1.36e-7 |
| LB (Amygdala) | R | PD>CT | +5.74 | 7.8e-7 | +0.156 | 0.502 | toward PD | +1.85e-7 | 1.71e-7 |
| **Other (n=20)** |  |  |  |  |  |  |  |  |  |
| Area 2 (Postcentral Sulcus) | L | PD>CT | +7.10 | 7.6e-9 | +0.413 | 0.245 | toward PD | +5.11e-7 | 5.11e-7 |
| Area 3b (Postcentral Gyrus) | L | PD>CT | +7.27 | 6.2e-9 | +0.397 | 0.338 | toward PD | +5.60e-7 | 4.03e-7 |
| Area OTS1 (Occipito-Temporal Sulcus) | L | PD>CT | +6.33 | 1.6e-7 | +0.876 | 0.089 | toward CT | -8.38e-7 | 4.92e-7 |
| Area Op1 (Parietal Operculum) | L | PD>CT | +7.56 | 2.7e-9 | +0.423 | 0.556 | toward PD | +2.08e-6 | 1.06e-6 |
| Area PFcm (Inferior Parietal Lobule) | L | PD>CT | +7.27 | 1.2e-8 | +0.633 | 0.301 | toward PD | +1.18e-6 | 5.78e-7 |
| Area STS2 (Superior Temporal Sulcus) | L | PD>CT | +6.51 | 9.6e-8 | +0.775 | 0.355 | toward PD | +2.26e-7 | 1.65e-7 |
| Area Te 1.0 (Heschl’s Gyrus) | L | PD>CT | +7.60 | 2.7e-9 | +0.481 | 0.624 | toward PD | +1.89e-6 | 1.59e-6 |
| Area Te 1.2 (Heschl’s Gyrus) | L | PD>CT | +7.08 | 1.7e-8 | +0.483 | 0.597 | toward PD | +1.94e-6 | 1.45e-6 |
| Area TeI (Superior Temporal Gyrus) | L | PD>CT | +7.16 | 1.4e-8 | +0.539 | 0.605 | toward PD | +2.79e-6 | 1.30e-6 |
| Area Op1 (Parietal Operculum) | R | CT>PD | +6.11 | 8.4e-8 | -0.075 | 0.371 | toward PD | +2.36e-6 | 1.69e-6 |
| Area Op3 (Parietal Operculum) | R | CT>PD | +2.55 | 0.015 | -0.313 | 0.405 | toward PD | +1.65e-6 | 1.10e-6 |
| Area PFcm (Inferior Parietal Lobule) | R | CT>PD | +5.75 | 2.7e-7 | -0.160 | 0.411 | toward PD | +1.73e-6 | 1.35e-6 |
| Area Ph2 (Parahippocampal Gyrus) | R | PD>CT | +6.35 | 1.0e-7 | +0.203 | 0.528 | toward PD | +5.17e-7 | 5.02e-7 |
| Area Te 1.0 (Heschl’s Gyrus) | R | PD>CT | +6.54 | 5.5e-8 | +0.249 | 0.550 | toward PD | +2.44e-6 | 1.66e-6 |
| Area Te 1.2 (Heschl’s Gyrus) | R | PD>CT | +5.91 | 3.6e-7 | +0.245 | 0.583 | toward PD | +3.38e-6 | 2.47e-6 |
| Area TeI (Superior Temporal Gyrus) | R | PD>CT | +6.17 | 1.5e-7 | +0.209 | 0.637 | toward PD | +1.96e-6 | 2.15e-6 |
| Area hOc4la (Lateral Occipital Suclus) | R | PD>CT | +6.14 | 2.2e-7 | +0.141 | 0.408 | toward PD | +1.98e-7 | 1.71e-7 |
| Area hOc5 (Lateral Occipital Sulcus) | R | PD>CT | +6.10 | 2.3e-7 | +0.222 | 0.254 | toward PD | +1.87e-7 | 1.72e-7 |
| Area hPO1 (Parieto-Occipital Sulcus) | R | CT>PD | +4.59 | 2.0e-5 | -0.181 | 0.543 | toward PD | +3.74e-7 | 1.44e-7 |
| Frontal-to-Occipital Gap | R | CT>PD | +4.79 | 8.8e-6 | -0.329 | 0.165 | toward PD | +1.50e-7 | 7.11e-8 |

### Pairwise Correlation & Functional Networks

Calculating pairwise regional saliency correlations revealed a different profile of underlying connectivity compared to raw saliency across subjects. General correlation was visualized via area specific node degree, which was the sum of the Spearman’s rho coefficients for significant correlations with other areas. We defined significance in this context to mean a pairwise correlation was significant in the majority of folds (pFDR < 0.05). Node degree gave a coarse picture of a region’s centrality without biasing towards regions that form few strong connections or regions that form many weak connections. Despite lower raw saliency scores, subcortical regions such as the SNc and other ROIs in the basal ganglia had higher node degrees especially as measured by GradCAM. Subcortical node degree was more left hemisphere biased for IG. Conversely, insular parcellations had higher raw saliency in both hemispheres but lower node degrees for both IG and GradCAM, suggesting the models observed more isolated and less network-level change in those regions (Fig 9).

**Figure 9:**
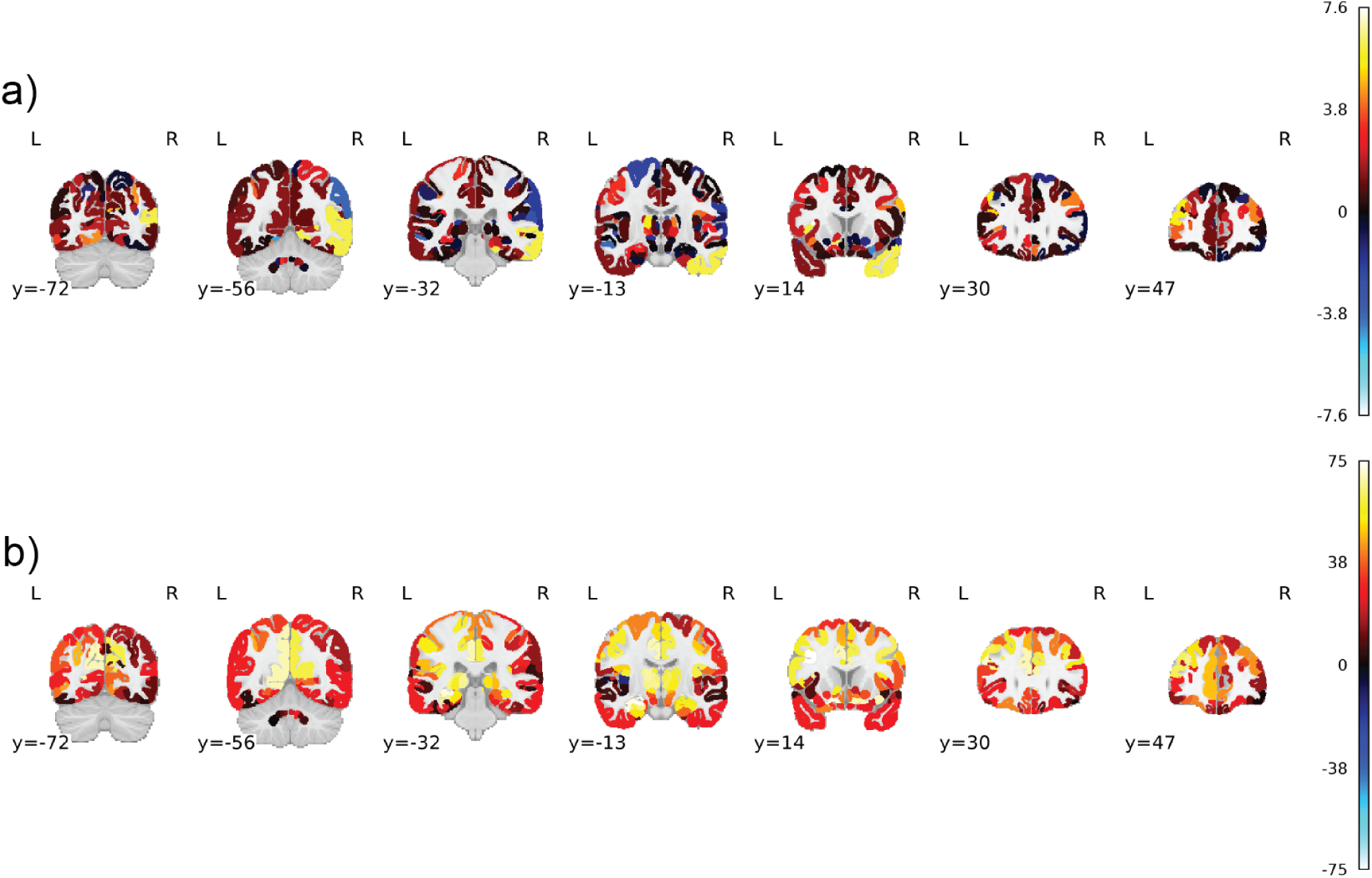
Coronal slice view of region node degree scoring for both GradCAM and Integrated Gradients (IG). A given area’s node degree is the sum of the mean Spearman’s coefficients across folds from significant correlations with all other regions. Panel a shows node degree per atlas region for IG, where panel b shows node degree for GradCAM. A pairwise correlation between two areas was only considered significant if three of five folds saw it as significant and the same sign (pFDR < 0.05). Greater node degree indicates an area’s saliency is either strongly correlated with the saliencies of other areas or that the area forms many significant correlations. L indicates left hemisphere, while R indicates right hemisphere.

While node degree provided a succinct metric to measure the tendency of an area to associate in networks of saliency brain-wide, it did not provide direct information on exactly which pairs of areas were highly correlated. To frame this analysis, we organized areas into canonical functional networks to see if saliency scores from areas within these networks tended to be stronger correlated with one another (within-functional network), or if our pairwise saliency correlations appeared to be agnostic to these functional connections (between-functional network) ^40–42^.

Functional network analysis revealed stronger correlations among areas within functional networks relative to their correlations with brain areas in different functional networks (Fig 10). Areas within the nigrostriatal pathway, consisting of the SNc, basal ganglia (including the striatum), and subthalamic nucleus (STN), correlated the strongest within-network and had the largest difference for within vs between network correlation for both GradCAM and IG (GradCAM average coefficient within-network: 0.56, between: 0.30, region-label permutation p < 0.0001; IG average coefficient within-network: 0.33, between 0.078, permutation p = 0.1102). It should be noted that IG’s voxel resolution makes it the better choice for this analysis as GradCAM’s 12x12x12 resolution is low and leads to higher overall correlation among adjacent areas. The region-label permutation test was also more biased for networks with smaller numbers of regions like the nigrostriatal (n=14), possibly explaining the lack of significance for IG despite the large effect size. Notably, despite their comparatively high degree of correlation, the nigrostriatal and limbic networks/pathways were lower in raw saliency for both CCGCAM and IG (Fig 11). Overall, pairwise correlation distinguished functional networks in a way raw saliency did not.

**Figure 10:**
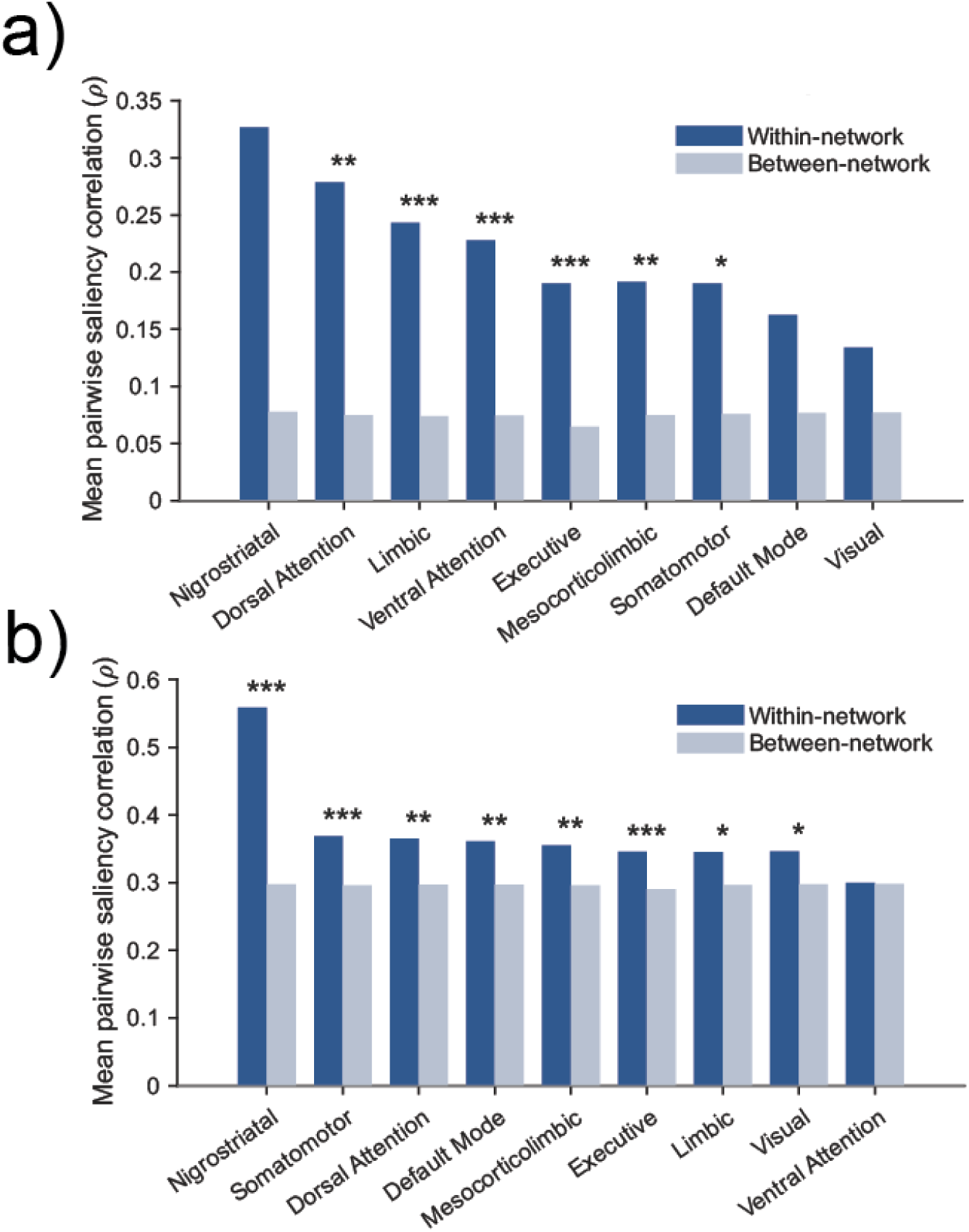
Functional networks are distinguished by pairwise saliency correlation for both GradCAM (GCAM) and Integrated Gradients (IG). Panel a shows within-network vs between-network correlation for IG, whereas panel b compares correlation for GradCAM. Bars show average correlation coefficient for significant correlations for regions within the same functional group (dark blue) versus average correlation coefficient for significant correlations for regions between different functional groups (light blue). Functional networks generally showed a higher average correlation coefficient among themselves than with all brain areas, though not every network reached significance in the region-label permutation test (see main text). Significance is labeled by asterisk as follows: *: <0.05, **: 0.01, ***: 0.001. Because pairwise correlations violate independence assumptions, statistical comparison was measured via random label permutations. Only fold-robust (3/5 folds pFDR< 0.05 and direction of correlation aligned) pairwise correlations were included in the analysis. A complete list of functional area segmentations is available in Supplemental Data 3.

**Figure 11:**
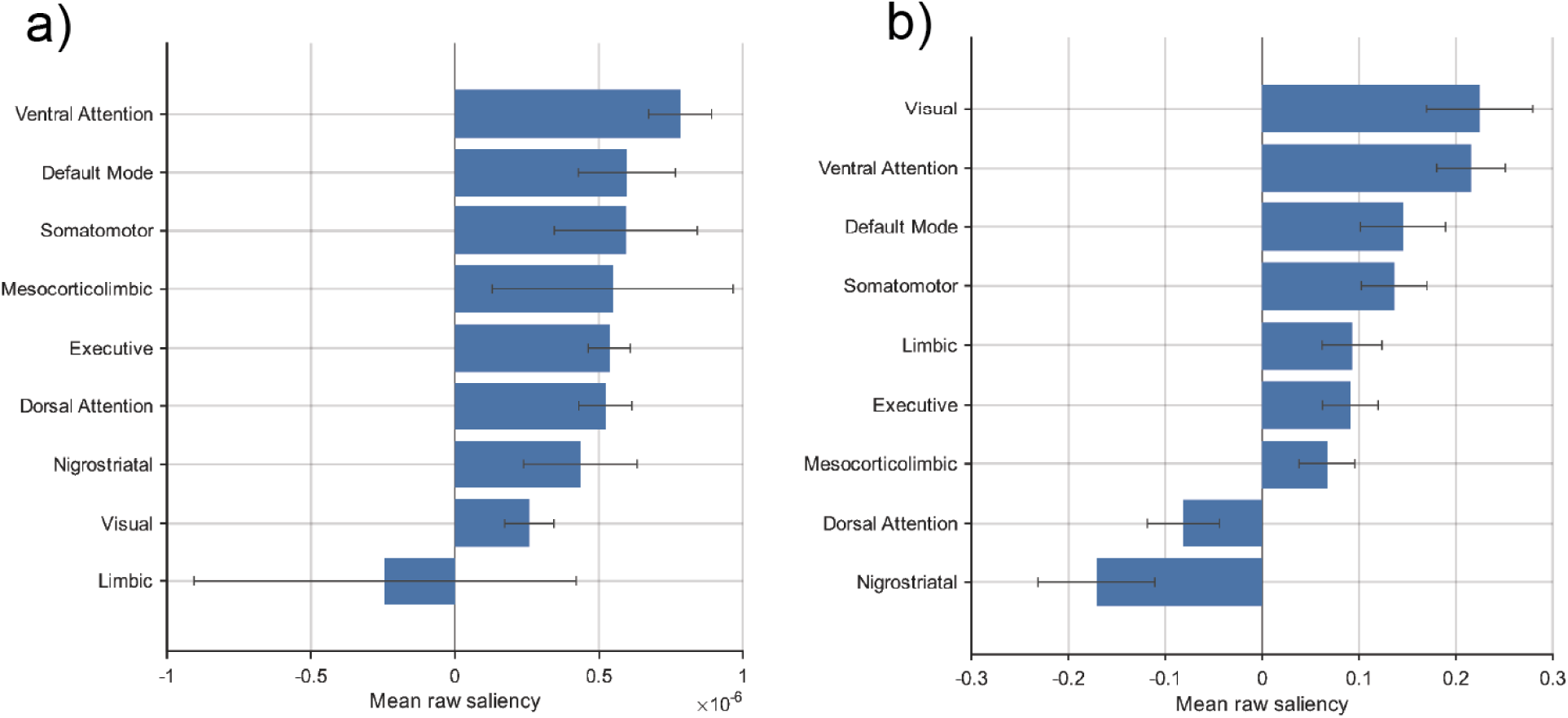
Functional network average saliency for both cohort contrastive GradCAM (CCGCAM) and Integrated Gradients (IG). Panel a shows IG average saliency for functional networks, where panel b shows GradCAM average saliency for functional networks. Mean raw saliency was taken by averaging robust saliency values for regions within different functional groups for both IG and CCGCAM. Robust values were determined by if all five folds agreed in sign of saliency for IG and if all five folds agreed in sign and were significantly different between PD and control cohorts for CCGCAM. Error bars show the standard error of mean for each functional network.

### Associations with Subject Clinical Metrics

Before examining correlations between clinical measures and our saliency data, we wished to understand any clinical-clinical relationships that existed within our holdout testing set and our 5-fold cross validation dataset. We tested scan level clinical-clinical correlation between UPDRS P3 total, MoCA total, LEDD, and years onset since symptoms. No clinical-clinical correlations were significant after FDR for our holdout testing set (likely due to the smaller sample size), but several relationships were observed in the cross-validation dataset used to train and validate the networks (Table 4). Notably, motor symptom progression was more strongly correlated with years since symptom onset in the unmedicated group relative to all PD subjects (ρ = 0.49, pFDR< 0.001 versus ρ = 0.37, pFDR< 0.001). There was no such correlation in the medicated group, suggesting dopamine replacement therapy interacted with the temporal trajectory of motor decline across subjects. Somewhat paradoxically, greater LEDD correlated with greater UPDRS P3 score, meaning subjects with more severe motor symptoms were likely on higher doses of medication. Because of this interaction, both metrics likely index overall PD progression.

**Table 4:** Significant pairs of correlated clinical metrics from subjects within the cross-validation dataset. Only correlations surviving FDR correction (pFDR < 0.05) are shown (None survived FDR for medicate subgroup). We performed all correlation as a scan level because scans from different timepoints from the same subject varied in clinical metrics. Scans were divided into medication subgroups (unmedicated: LEDD = 0, medication: LEDD > 0) for individual analysis to understand how medication status may influence clinical trajectories. Correlations for on-medication scans in the cross-validation and all groups within the holdout testing set were also calculated but did not reach significance and are not shown. Subjects with missing data were withheld from analysis. LEDD: Levadopa equivalent daily dose, UPDRS-III: Unified Parkison’s Disease Rating Scale Part 3. See also: Supplemental Data 7 shows clinical metrics in unmedicated vs medicated subgroups in the holdout testing set.

| Clinical measure pair | Coefficient (Spearman) | pFDR | n (scans) |
|---|---|---|---|
| **Cross Validation Set** scans = 202 |  |  |  |
| LEDD (mg) × Yrs since symptom onset | 0.39 | pFDR < 0.001 | 197 |
| UPDRS-III total × Yrs since symptom onset | 0.37 | pFDR < 0.001 | 190 |
| UPDRS-III total × LEDD (mg) | 0.35 | pFDR = 0.002 | 194 |
| **Cross Validation, Off-Medication** scans = 132 |  |  |  |
| UPDRS-III total × Yrs since symptom onset | 0.49 | pFDR < 0.001 | 128 |

After establishing the relationship between different clinical metrics within our dataset, we investigated correlations between our CNN-derived regional saliency scores and clinical metrics to ground our abstract saliency results to concrete subject data. While we could have analyzed correlations down to an atlas area resolution, doing so would result in 414 correlations per clinical metric we wished to test. For this exploratory and hypothesis generating analysis, we pooled areas into lateralized lobes and functional networks. This pooling made correlation results less fragile and more robust across folds and was more compatible with the low spatial resolution of GradCAM. To maximize the sample size for correlation, we performed this analysis at scan level. For each scan, we averaged saliency scores over pooled lobes and functional networks and calculated correlation across scan pooled region saliency with each of the following clinical metrics taken from the nearest timepoint to the image: total MoCA score, total UPDRS P3 score, LEDD, and years since onset of symptoms. We observed the most robust correlations across many lobes and functional networks for both UPDRS P3 (Fig 12) and LEDD (Fig 13).

**Figure 12:**
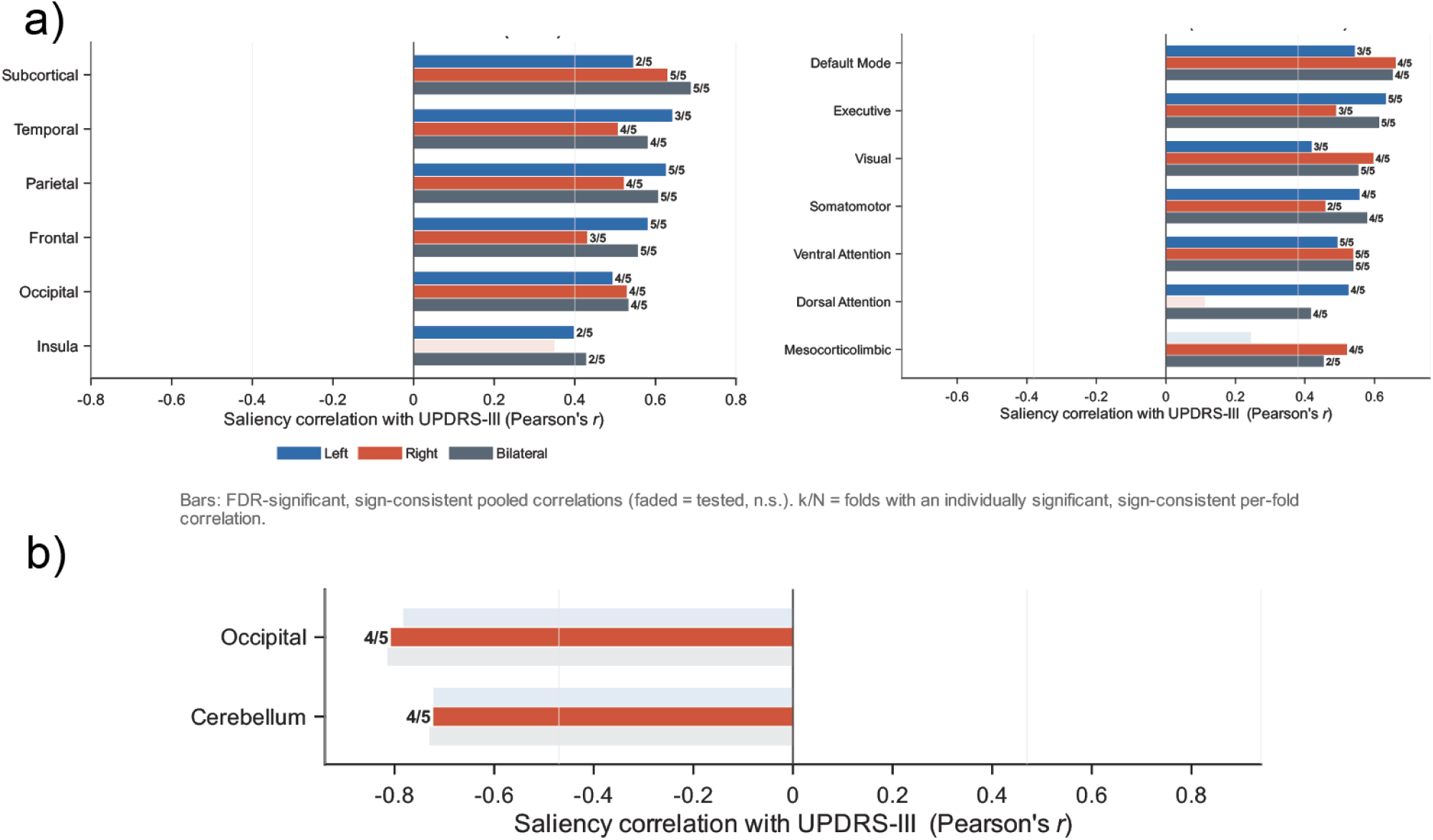
Correlations between functional network and lobe saliency and UPDRS P3 motor symptom measures for both Integrated Gradient (IG) and GradCAM. Panel a shows IG pooled region saliency scores correlated with UPDRS P3 scores, where panel b shows GradCAM saliency correlations. Different hemispheres are shown by different color bars (blue: left, orange: right, grey: both). Number of robust folds are shown next to bar plots, defined by the number of folds that had significant correlations (pFDR < 0.05) in the same direction. Correlations were calculated at scan level. Nonsignificant hemispheres are shown greyed out for lobes/functional networks with other significant hemispheres. If a lobe/functional network formed no significant correlation, it was omitted. Sample size varied between folds, so for exploratory visualization robustness criteria were relaxed to two or more folds seeing significant, same sign correlation (fold 1: n = 30, fold 2: n = 30, fold 3: n = 19, fold 4: n = 43, fold 5: n = 20).

**Figure 13:**
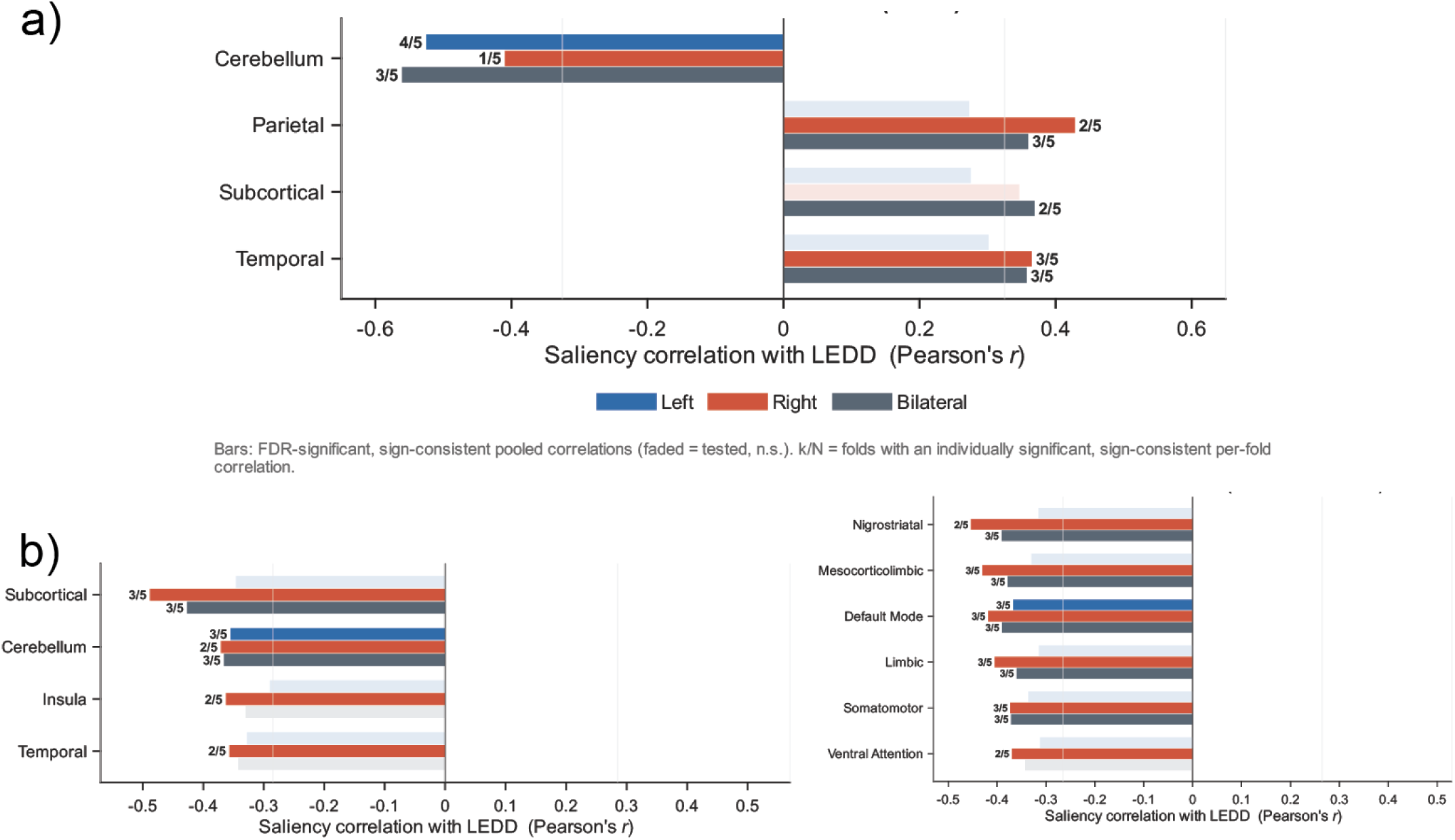
Correlations between functional network and lobe saliency and LEDD for both Integrated Gradient (IG) and GradCAM. Panel a shows IG pooled region saliency scores correlated with LEDD, where panel b shows GradCAM saliency correlations. Different hemispheres are shown by different color bars (blue: left, orange: right, grey: both). Number of robust folds are shown next to bar plots, defined by the number of folds that had significant correlations (pFDR < 0.05) in the same direction. Correlations were calculated at scan level. Nonsignificant hemispheres are shown greyed out for lobes/functional networks with other significant hemispheres. If a lobe/functional network formed no significant correlation, it was omitted. Sample size varied between folds, so for exploratory visualization robustness criteria were relaxed to two or more folds seeing significant, same sign correlation (fold 1: n = 30, fold 2: n = 30, fold 3: n = 19, fold 4: n = 43, fold 5: n = 20).

UPDRS P3 correlated with a variety of lobes and functional networks for IG. Correlation between motor symptoms and IG was uniformly positive, indicating that higher IG saliency was associated with worse motor symptoms across all pooled regions. Bilateral subcortical areas had the largest positive effect (r = 0.69, 5/5 folds robust), consistent with the subcortical basis of dopaminergic pathology thought to underpin motor deficits in PD. The same effect is absent from the nigrostriatal pathway, suggesting the subcortical correlation was driven largely by thalamic and amygdalar saliency. Positive correlation with IG was largely distributed across many lobes (including frontal, temporal, and parietal) and functional networks (executive, somatomotor, default mode). This could indicate that motor symptoms relate to system-wide brain change, and thus to PD pathology as a whole, rather than to highly localized change. In summary, IG tracked motor symptom progression across a wide variety of lobes and functional networks. While GradCAM formed few significant correlations with UPDRS P3 overall, the correlations between the occipital lobe and cerebellum were negative, opposite the direction of the positive correlations observed from IG. Despite the diverging sign, this suggested that GradCAM was tracking a similar progression related signal as IG.

Like UPDRS P3, LEDD was significantly correlated with a variety of lobes and functional networks but primarily in GradCAM, with less correlation in IG. Correlation between GradCAM and LEDD was uniformly negative, indicating that GradCAM saliency decreased across many pooled regions as medication dosage increased. This suggests medication status was associated with reduced GradCAM sensitivity to PD across a wide area of pooled regions. This was contrasted by IG’s positive correlation with LEDD in the temporal, subcortical, and parietal lobes. Within each of GradCAM and IG, the sign of correlation with UPDRS P3 and with LEDD mostly agreed across similar pooled regions, reflecting that both methods may see a relationship between UPDRS P3 and LEDD. LEDD and UPDRS P3 were correlated in our cross-validation set (Table 4) and so it’s likely that both measures were entangled with overall PD severity. IG and GradCAM both agreed in sign of correlation in the cerebellum only, demonstrating a method-robust decreasing signal with more progression. The only correlation between saliency and a metric other than LEDD or UPDRS P3 was between the left and bilateral cerebellum and years since symptom onset for IG (r = -0.77, pFDR < 0.001, 4/5 folds robust; r = -0.7, pFDR < 0.001, 3/5 folds robust respectively). Altogether, this suggests cerebellar saliency may be highest in very early PD, before subjects experience significant motor symptoms or receive treatment.

## Discussion

Through patterns in raw saliency and its correlations with motor scores, we find that atrophy could have been one of multiple anatomical features contributing to regional saliency scores generated both through cohort contrastive GradCAM (CCGCAM) and Integrated Gradients (IG). Raw saliency was left lateralized for both methods, corroborating other studies’ findings that atrophy largely affects the left hemisphere in early PD ^15,45^. The pattern of saliency seemed to show a threshold of pathology at which the network could make accurate predictions. This threshold appeared to start at Braak Stages 3 and 4 as evidenced through robust saliency in the ventral striatum and amygdala. More globally, average saliency across subjects best reflected Braak Stages 5 and 6 where pathology spreads to the neocortex ^16^. Furthermore, pairwise correlation between area saliency scores measured by IG and GradCAM linked areas known to be functionally connected. Correlations within functional networks were greatest within the nigrostriatal pathway for both IG and GradCAM. Despite being the most correlated, the difference between within-network and between-network correlation for nigrostriatal areas reached permutation significance only for GradCAM. This is consistent with the CNN classifier tracing PD-related anatomical patterns collectively across connected areas, and with theories of network-level effects of PD neuropathology within dopaminergic pathways. While subcortical saliency had the highest correlation with motor symptoms for IG, correlation with motor symptoms was widespread. It appears saliency tracked PD progression primarily along the axis of motor symptoms rather than cognitive performance or duration, as years since first symptoms correlated only with the cerebellum for IG and MoCA formed no correlations. The lack of correlation with metrics other than UPDRS P3 suggests anatomical differences observable brain wide in T1w MRI may be more connected to motor deficits than other symptoms or clinical features such as disease duration. Increasing medication use appeared to reflect PD progression: LEDD correlated with both time since first symptoms and motor symptoms and showed a saliency-correlation profile similar to that of UPDRS P3.

We successfully extracted comprehensive quantifications of area-specific saliency data from a “black box” 3D CNN model. We next sought to understand what the saliency output means and why it could be informative. Fundamentally, our area-specific saliency scores provided voxel-wise (IG) and coarse (CCGCAM) quantifications of the regional discriminative importance the network assigned when separating control from PD. They did not, however, explicitly define what those differences were in anatomical terms beyond the fact that they were consequential to the network’s classification of a subject. A given area may have been salient for PD because it showed atrophy relative to control, but this was an inference that we clarified by comparing saliency with cortical thickness findings from literature and through establishing a connection between saliency and subject attributes associated with atrophy, like cognitive scores and motor performance assessments.

We hypothesized that gray matter atrophy could have been an anatomically significant signal that was picked up by our saliency measurements. We observed that younger controls were correctly classified at a higher rate than older controls (Supplemental Data 6). It is possible the model conflated age-related thinning in control images with PD pathology, which would suggest it relied on atrophy as a general feature of PD. Comparison with cortical thickness literature in PD further supported the atrophy hypothesis but also revealed its limits. Areas with highest raw saliency in our data mirrored areas with highest gray matter volume reduction in early PD reported by Claassen et al. 2016. This study measured differences in atrophy between a late and early group of PD subjects, revealing a left hemisphere-biased profile of atrophy in the frontal lobe and insula as well as modest bilateral atrophy in the operculum and temporal regions in early PD, which progressively increased bilaterally and toward posterior areas in later PD subjects (over 10 years). Our dataset was composed entirely of early subjects, and our saliency scores reflect this in comparison with atrophy profiles from literature ^15,45^. Saliency was left lateralized in the frontal lobe especially for CCGCAM and, when combined with insular and temporal regions, formed a left hemisphere biased robust cluster (Fig 7, 8). Both IG and CCGCAM had higher saliency clusters of regions around the operculum in both the right and left hemispheres. This result was also consistent with the profile of early PD atrophy in the literature ^15,45^. Our saliency data reflected brain-wide correlations with motor symptoms (Fig 12). This is consistent with findings that atrophy in a diverse set of regions correlates with motor symptoms ^46,47^. In summary, saliency scores are broadly consistent with reported atrophy profiles and with their connection to motor symptoms in PD, supporting the hypothesis that atrophy could have been a robust feature contributing to models’ saliency scores.

While our raw saliency profiles were largely consistent with that of gray matter atrophy in early PD and with the nature of gray matter atrophy’s connection to motor symptoms, saliency was not as consistent with respect to gray matter atrophy’s connection with cognitive decline. Longitudinal cortical thickness measurements made with tools such as FreeSurfer have demonstrated an association between atrophy in PD and declining cognitive scores ^13^. We did not observe any significant correlation with cognitive scores in our dataset, but this could be due to the nature of cognitive decline in PD. Firstly, cognitive decline is not as universal as motor difficulties among people with PD, especially early on in progression. Estimates place the number of subjects experiencing cognitive decline within the first 10 years of PD at 20%, increasing to 80% among subjects who had had PD for more than 10 years ^48^. Our dataset was composed entirely of subjects who’d had PD symptoms for 10 years or less, meaning cognitive decline was not as well represented in our dataset. In fact, among our holdout testing subjects, cognitive scores were not significantly lower in the PD cohort than in controls (Table 1) and were relatively high for PD subjects in the entire dataset (MoCA PD mean: 27.1 SD: 2.7, 25 or lower is mild cognitive impairment). We may have observed a robust correlation between cognitive decline with the inclusion of more subjects who’d experienced greater cognitive decline, which would necessitate many longer-term cases of PD. Our dataset was also cross-sectional and did not have the same longitudinal perspective as studies that found a correlation between atrophy and cognitive symptoms. Cognitive decline in PD may involve subsets of diffused cortical areas with variability between subjects, which in conjunction with the underrepresentation of more severe cognitive impairment in the dataset, could have made it difficult to observe a regular profile across subjects ^49^. It should also be noted that, apart from the cerebellum, no correlations with years since onset survived FDR correction. Like with cognitive symptoms, this could be explained by the limited diversity of durations in our entirely early PD dataset. In summary, atrophy appears to be an anatomical feature captured by saliency. It is unlikely to have been the only feature captured by saliency, however. Saliency is a high dimensional metric and could have captured subtle structural changes connected to changes in connectivity, other pathological features separate from atrophy, or spurious correlations learned by the models. While this can weaken the anatomical interpretation of saliency, it also provides a strong platform for a diverse set of analyses. Where a measure like cortical thickness will ignore changes in white matter hyperintensity, for example, our 3D CNN models can capture those changes if they help the network distinguish between classes. This opens multiple possibilities for different ways to segment or atlas saliency maps. As white matter atlases improve and become more popular, saliency could be a generalizable tool to apply to more than only gray matter segmentation templates and atlases ^50^.

A well-trained CNN could be sensitive to a variety of features indicative of PD, and saliency can reveal which of those features are present for a given individual. To respect the heterogeneity that existed within our data, we analyzed how different areas’ saliencies change with one another independent of the magnitude of saliency through pairwise area correlation.

We observed that subcortical areas were more associated brain-wide via rank-based pairwise correlation than they were high saliency, as summarized by node degree (Fig 9). We then established specific patterns of correlation using functional networks as a frame, with an eye on subcortical areas participating in dopaminergic pathways heavily implicated in PD pathology. This also provided insight into whether our 3D CNN could pick up trends in structural images rooted in underlying functional connectivity. Generally, saliencies of areas within functional networks associated with one another considerably more than they associated with other brain areas in different functional networks, revealing that saliency changes tend to occur over groups of areas that canonical parcellations assign to the same functional network (Fig 10). The differing magnitudes between within-network and between-network correlation varied in a way that tracked networks implicated in PD. The heavily implicated nigrostriatal pathway, which includes the SNc, striatum, and STN, showed the greatest differential in within-network vs between-network correlation (Fig 10; 0.252 difference in Spearman coefficient for IG). This correlation structure distinguished the pathway, which was otherwise low in raw saliency (Fig 11). These findings suggest that saliency captured second order relationships between areas that were not visible in raw saliency alone. In addition, the correlation structure of saliency aligned with canonical functional network boundaries drawn from the literature, though we did not measure functional connectivity in these subjects. That the largest such differential fell on the nigrostriatal pathway for both GradCAM and IG is consistent with its established involvement in PD pathology. More generally, our model captured network-aligned saliency associations across both attribution methods, raising the possibility that 3D CNNs recover correlates of functional organization from structural data alone.

Braak staging helps frame a neurobiologically grounded threshold of pathology at which the models could correctly classify subjects. Our method-robust saliency data highlights an array of regions in prefrontal, parietal, and temporal cortex, as well as subcortical structures (Table 3). This pattern is consistent with the model settling at a pathology threshold around Braak Stages 3 or 4, where pathology spreads to subcortical areas such as the SNc and amygdala and begins to impact temporal regions ^16^. Robust neocortical saliency captured elements of pathology described in Braak Stages 5 and 6, where changes are visible in a wider swath of areas across the brain.

In summary, Braak Staging affords an interpretation of our saliency map as the threshold of pathology preceding the models’ ability to make an accurate prediction. The relatively moderate classification accuracy of the model makes sense given the early PD makeup of our dataset and the fact the pathology map reflects more progressed Braak Stages of PD pathology. It is notable that, despite the apparent connection of saliency to later Braak Stages, the cerebellum was the only pooled region whose saliency correlated with years since first symptoms after FDR correction. The lack of connection could be attributed to the heavy early bias of our dataset and the moderate sample size for this analysis.

Saliency formed correlations with both UPDRS P3 and LEDD across a broad set of pooled regions for both IG and GradCAM, while years since onset correlated with the cerebellum for IG only and MoCA formed no significant correlations for any method. Within each saliency method, the sign of correlation was generally maintained across both LEDD and UPDRS P3. Furthermore, the correlated pooled regions generally did not diverge between LEDD and UPDRS P3. Combining these findings with the observation that UPDRS P3 and LEDD were correlated in the cross-validation portion of our dataset, it is likely that UPDRS P3 and LEDD were entangled to give a general measure of progression instead of showing symptom specific anatomical changes.

The cerebellum was the only pooled region in which IG and GradCAM agreed in sign and significance for both LEDD and UPDRS P3. It was also the only pooled region that formed any connection with years since symptom onset, having a negative correlation with IG bilaterally and in the left hemisphere. Because all these correlations were negative, this method-robust signal suggests cerebellar saliency was highest earlier in the disease course and in subjects with milder motor symptoms and lower medication burden. It has been found that cerebellar gray matter is reduced among several nuclei in the cerebellum for early PD, and that the functional connectivity between them is negatively linked with posture scores ^51^. Our result is consistent with that work, though we measured saliency rather than functional connectivity and so cannot speak to the connectivity directly. Regardless, these results are consistent with the cerebellum undergoing change early in PD progression, possibly reflecting a compensatory response as motor function begins to deteriorate.

Across different levels of analysis, we observed a varying level of agreement between GradCAM and IG. Raw saliency signals highlighted similar regions between the two methods. Pairwise regional correlation was much greater in magnitude for GradCAM across all regions, including those within the same functional network, but the pattern of greater within-network correlation held for both methods. The methods agreed on which clinical metrics they correlated with, but the regions involved were largely non-overlapping and the correlations were opposite in sign with the exception of the cerebellum.

Although one might expect the two methods to overlap completely, they diverge mechanistically and therefore give different information about PD features on a per-subject basis. GradCAM shows the spatial attributions of abstract features the model learns, coarsely showing where the discriminative features were localized. GradCAM samples activations from deep convolutional layers and produces a low resolution, fully positive map (12×12×12 in our application) showing sensitivity to where class-supporting features lie. This is in contrast with IG, which tests the individual contribution of specific voxels to a classification outcome. It outputs a map of how much each individual voxel pushed the network toward or away from classification relative to a baseline volume. IG is signed, has voxel resolution, and is complete (the attributions sum to the class score difference between image and baseline).

The similarities and divergences between the two attribution methods in our results tracked their mechanistic differences. Raw saliencies between the methods may have been more similar because the volatile signed effects of IG averaged out across subjects and converged on GradCAM’s coarser profile. Similarly, GradCAM’s coarse resolution provides a trivial explanation for its higher pairwise correlation between areas relative to IG. The opposing signs of their UPDRS P3 and LEDD correlations (positive for Integrated Gradients, negative for GradCAM) may reflect that the two methods measure different quantities. GradCAM’s sensitivity itself is abstract and could have been measuring the intactness of certain areas, leading to the magnitude of GradCAM’s signal being inverse with progression. Despite negative directionality, there was still sensitivity to these regions which correlated to progression. Conversely, IG’s completeness explains positive correlations with severity of progression. The sum of IG’s attributions across all voxels equals the difference in logit between the specific PD scan and control baseline. If more progressed PD cases produce larger logits for the PD class, the difference between the PD logit and the control baseline logit would also be greater. A greater logit difference would in turn yield a greater share of positive attributions in more severe cases, which would explain the widespread positive correlation. We did not test this directly.

Our model performed with moderate accuracy on the difficult task of PD classification from T1w MRI. Several factors likely contributed to the models’ difficulty in identifying early PD. Firstly, we applied a rigorous site holdout scheme to each set of subjects used to validate or test the networks, ensuring all testing and evaluation was performed on data from unseen sites. In the literature, models evaluated with explicit site stratification give less optimistic generalizability estimates. The site stratified model from Camacho et al., 2023 performed with 79.3% accuracy (trained with 2,041 subjects), while the model from Chakraborty et al. 2020 reported 92.75% without comparable site controls on a smaller dataset of 406 MRIs. A site stratified scheme was essential for our study, as our dataset included images from 39 different imaging sites, many of which contributed subjects of only one cohort. We used heavy augmentation to increase regularization and overcome some site bias; however, biased folds were difficult to avoid without more complex harmonization through preprocessing. We sought to prioritize honest evaluation over highly optimized accuracy. This rigid approach to evaluation exacerbated the impact of our limited dataset (141 PD and 126 control subjects, of whom 30 PD and 29 controls were split off for holdout testing). We selected the most well-balanced sites for testing, leaving many of the sites in the cross-validation set heavily biased, which likely impacted the models’ training. This influence is evidenced in our saliency data by heavy per-fold idiosyncrasies (Fig 5, 6): data from different imaging sites may have led each fold to learn different features. We applied strict fold robustness criteria to restrict the influence of these idiosyncrasies and to make our saliency analysis reproducible across folds, but these idiosyncrasies likely remained influential in limiting the networks’ ability to generalize to new imaging sites.

Secondly, despite our interfold robustness screening, the ensemble model appeared to conflate age with pathology as evidenced by more difficulty classifying older control subjects. This weakens our interpretation of saliency as strictly pathology-correspondent, as the models may have highlighted general age-related features alongside PD-related ones. Finally, our dataset was composed of subjects very early in the PD time course who may naturally show limited anatomical evidence of PD in T1w MRI. It should be noted that the metric we used to summarize progression (years since onset of symptoms) reflected a longer time course than the PPMI reported years of disease duration (mean years since symptom onset in test set: 3.8, mean disease duration 0.6). Especially in particularly early cases, it is unlikely our dataset of 141 PD and 126 control MRI subjects captured a complete profile of the heterogeneous PD-associated structural features available in T1w MRI. The model would have benefited from more MRI data from PPMI to further increase performance and confidence in evaluation on the difficult task of classifying early PD.

Our preprocessing may also have introduced small errors. The atlasing approach relied on the linear registration of all subjects to the same anatomical space to align Julich Brain parcellations with subject data. It was therefore impossible for the atlas to be perfectly accurate subject to subject; this was partly mitigated by probabilistic parcellations, in which each voxel carries a probability of belonging to an atlas area rather than a binary assignment. This helped normalize the influence of anatomical variability between subjects regardless of registration. Nevertheless, there were still small errors with intersubject atlas alignment. Innovative deep-learning-based per subject segmentation techniques could be employed to increase atlas alignment accuracy, provided error is considered and validated ^52^.

To align the atlas, all MRIs had to be linearly registered to MNI152 space. This could bias the representation of brain regions across subjects, as all volumes were warped during registration to fit the template anatomical space ^53^. This could be especially impactful when registering MRIs of subjects with PD pathology, as pathological change may alter how those areas register relative to control brains ^54^. PD-based anatomical spaces such as PD25 exist and could have been the template chosen for spatial normalization, but they are not natively compatible with the Julich Brain Atlas. While the atlas could theoretically be registered to another anatomical space, this process has not been validated, and controls would also have needed to be registered to a pathological anatomical space.

In this study, we propose an ensemble of 3D CNN models capable of classifying PD in T1w MRIs from unseen imaging sites with 67.8% accuracy. We introduce a multi-method, multi-scale, and robust explainability pipeline to interpret the anatomical signals used by these traditionally opaque models in a way that can be thoroughly quantitatively analyzed. We analyzed the diverse set of features this metric contains, showing that it can be connected to cortical thickness and to pathology thresholds characterized by Braak staging, and that its correlation structure aligns with canonical functional networks derived from structural data alone. We also connected saliency scores to general PD progression through motor symptoms and medication, with subcortical regions’ saliency being the most correlated to motor symptoms. This study should be repeated on a larger cohort of PD subjects from PPMI, with cortical thickness analysis performed on the same dataset used to train the 3D CNN model to allow for direct connection between atrophy and saliency. This study could also be replicated for detection of any variety of pathologies where large quantities of MRI data are available using any 3D CNN architecture, as both GradCAM and Integrated Gradients are architecture-agnostic ^34,36^. Within PPMI data alone, prodromal MRIs could be added to the dataset to establish anatomical differences that exist between control, prodromal, and PD. More work can also be done to clarify features captured by saliency. Saliency could also be compared with any subject metric, not just the few used in this study (UPDRS P3, MoCA, time since symptom onset, LEDD). Saliency is a complex metric that is influenced by a variety of features and information from training MRIs. Linkages and associations can be investigated and clarified by examining how saliency connects to a multitude of metrics in medical imaging analysis.

## Data Availability

Data used in the preparation of this article was obtained on 2026-01-05 from the Parkinson’s Progression Markers Initiative (PPMI) database (www.ppmi-info.org/acces-dataspecimens/download-data), RRID:SCR_006431. For up-to-date information on the study, visit www.ppmi-info.org.

Statistical atlas parcellations were retrieved on 2025-06-17 from the Julich Brain Cytoarchitectonic Atlas v3.1 using the Siibra Python Package ^39^.

## Code Availability

Code is available upon request to the corresponding author.

## Acknowledgements

PPMI – a public-private partnership – is funded by the Michael J. Fox Foundation for Parkinson’s Research and funding partners, including 4D Pharma, Abbvie, AcureX, Allergan, Amathus Therapeutics, Aligning Science Across Parkinson’s, AskBio, Avid Radiopharmaceuticals, BIAL, BioArctic, Biogen, Biohaven, BioLegend, BlueRock Therapeutics, Bristol-Myers Squibb, Calico Labs, Capsida Biotherapeutics, Celgene, Cerevel Therapeutics, Coave Therapeutics, DaCapo Brainscience, Denali, Edmond J. Safra Foundation, Eli Lilly, Gain Therapeutics, GE HealthCare, Genentech, GSK, Golub Capital, Handl Therapeutics, Insitro, Jazz Pharmaceuticals, Johnson & Johnson Innovative Medicine, Lundbeck, Merck, Meso Scale Discovery, Mission Therapeutics, Neurocrine Biosciences, Neuron23, Neuropore, Pfizer, Piramal, Prevail Therapeutics, Roche, Sanofi, Servier, Sun Pharma Advanced Research Company, Takeda, Teva, UCB, Vanqua Bio, Verily, Voyager v. 25MAR2024 Therapeutics, the Weston Family Foundation and Yumanity Therapeutics.

## Author Contributions

Conceptualization: EM and AY; Experimental procedures: EM; Analysis: EM, BZ, AY; Writing – original draft: EM, AY; Writing – review & editing: EM, BZ, AY.

## References

1. de Lau, L. M. L. & Breteler, M. M. B. Epidemiology of Parkinson’s disease. Lancet Neurol. 5, 525–535 (2006).

2. de Rijk, M. C. et al. Prevalence of parkinsonism and Parkinson’s disease in Europe: the EUROPARKINSON Collaborative Study. European Community Concerted Action on the Epidemiology of Parkinson’s disease. Journal of Neurology, Neurosurgery & Psychiatry 62, 10 (1997).

3. Cramb, K. M. L., Beccano-Kelly, D., Cragg, S. J. & Wade-Martins, R. Impaired dopamine release in Parkinson’s disease. Brain 146, 3117–3132 (2023).

4. Bae, Y. J. et al. Imaging the Substantia Nigra in Parkinson Disease and Other Parkinsonian Syndromes. Radiology 300, 260–278 (2021).

5. Armstrong, M. J. & Okun, M. S. Diagnosis and Treatment of Parkinson Disease: A Review. JAMA 323, 548–560 (2020).

6. Mavridis, I. & Pyrgelis, E.-S. Nucleus accumbens atrophy in Parkinson’s disease (Mavridis’ atrophy): 10 years later. Am. J. Neurodegener. Dis. 11, 17–21 (2022).

7. Hanganu, A. et al. Mild cognitive impairment is linked with faster rate of cortical thinning in patients with Parkinson’s disease longitudinally. Brain 137, 1120–1129 (2014).

8. Kobayakawa, M., Tsuruya, N. & Kawamura, M. Decision-making performance in Parkinson’s disease correlates with lateral orbitofrontal volume. J. Neurol. Sci. 372, 232–238 (2017).

9. Adachi, M., Hosoya, T., Haku, T., Yamaguchi, K. & Kawanami, T. Evaluation of the Substantia Nigra in Patients with Parkinsonian Syndrome Accomplished Using Multishot Diffusion-Weighted MR Imaging. AJNR Am J Neuroradiol vol. 20 (1999).

10. Heim, B., Krismer, F., De Marzi, R. & Seppi, K. Magnetic resonance imaging for the diagnosis of Parkinson’s disease. J. Neural Transm. 124, 915–964 (2017).

11. Seifert, K. D. & Wiener, J. I. The Impact of DaTscan on the Diagnosis and Management of Movement Disorders: A Retrospective Study. Am J Neurodegener Dis vol. 2 www.AJND.us/ (2013).

12. Pagano, G., Niccolini, F. & Politis, M. Imaging in Parkinson’s disease. Clinical Medicine 16, 371–375 (2016).

13. Pletcher, C. et al. Cerebral cortical thickness and cognitive decline in Parkinson’s disease. Cereb. Cortex Commun. 4, tgac044 (2023).

14. Weintraub, D. et al. Neurodegeneration Across Stages of Cognitive Decline in Parkinson Disease. Arch. Neurol. 68, 1562–1568 (2011).

15. Claassen, D. O. et al. Cortical asymmetry in Parkinson’s disease: early susceptibility of the left hemisphere. Brain Behav. 6, e00573 (2016).

16. Braak, H. et al. Staging of brain pathology related to sporadic Parkinson’s disease. Neurobiol. Aging 24, 197–211 (2003).

17. Patel, S. B., Goh, V., Fitzgerald, J. J. & Antoniades, C. A. Multi-Cohort Development and Validation of 2D and 3D Deep Learning Models for MRI-Based Parkinson’s Disease Classification: Comparative Analysis of ConvKANs, CNNs, and GCNs. https://arxiv.org/abs/2407.17380 (2024).

18. Crespi, L., Loiacono, D. & Sartori, P. Are 3D better than 2D Convolutional Neural Networks for Medical Imaging Semantic Segmentation? in 2022 International Joint Conference on Neural Networks (IJCNN) 1–8 (2022). doi:10.1109/IJCNN55064.2022.9892850.

19. Chang, K. et al. Distributed deep learning networks among institutions for medical imaging. Journal of the American Medical Informatics Association 25, 945–954 (2018).

20. Marek, K. et al. The Parkinson’s progression markers initiative (PPMI) – establishing a PD biomarker cohort. Ann. Clin. Transl. Neurol. 5, 1460–1477 (2018).

21. Simuni, T. et al. Longitudinal Change of Clinical and Biological Measures in Early Parkinson’s Disease: Parkinson’s Progression Markers Initiative Cohort. Movement Disorders 33, 771–782 (2018).

22. Marek, K. et al. The Parkinson Progression Marker Initiative (PPMI). Prog. Neurobiol. 95, 629– 635 (2011).

23. Chakraborty, S., Aich, S. & Kim, H. C. Detection of Parkinson’s disease from 3T T1 weighted MRI scans using 3D convolutional neural network. Diagnostics 10, (2020).

24. Rustom, F., Moroze, E., Parva, P., Ogmen, H. & Yazdanbakhsh, A. Deep learning and transfer learning for brain tumor detection and classification. Biol. Methods Protoc. 9, bpae080 (2024).

25. Rosenbacke, R., Melhus, Å., McKee, M. & Stuckler, D. How Explainable Artificial Intelligence Can Increase or Decrease Clinicians’ Trust in AI Applications in Health Care: Systematic Review. JMIR AI vol. 3 Preprint at 10.2196/53207 (2024).

26. Rosen, A. F. G. et al. Quantitative assessment of structural image quality. Neuroimage 169, 407– 418 (2018).

27. Kim, H. E. et al. Transfer learning for medical image classification: a literature review. BMC Med. Imaging 22, 69 (2022).

28. Ebrahimi, A., Luo, S. & Initiative, A. D. N. Convolutional neural networks for Alzheimer’s disease detection on MRI images. Journal of Medical Imaging 8, 024503 (2021).

29. Glorot, X. & Bengio, Y. Understanding the Difficulty of Training Deep Feedforward Neural Networks. http://www.iro.umontreal. (2010).

30. He, K., Zhang, X., Ren, S. & Sun, J. Deep Residual Learning for Image Recognition. http://image-net.org/challenges/LSVRC/2015/ (2016).

31. Kushol, R., Parnianpour, P., Wilman, A. H., Kalra, S. & Yang, Y.-H. Effects of MRI scanner manufacturers in classification tasks with deep learning models. Sci. Rep. 13, 16791 (2023).

32. Avants, B. B. et al. Magnetic Resonance Imaging Data Phenotypes for the Parkinson’s Progression Markers Initiative. Preprint at 10.1101/2024.09.23.24313179 (2024).

33. Cirillo, M. D., Abramian, D. & Eklund, A. WHAT IS THE BEST DATA AUGMENTATION FOR 3D BRAIN TUMOR SEGMENTATION? in Proceedings - International Conference on Image Processing, ICIP vols 2021-September 36–40 (IEEE Computer Society, 2021).

34. Selvaraju, R. R. et al. Grad-CAM: Visual Explanations from Deep Networks via Gradient-Based Localization. in 2017 IEEE International Conference on Computer Vision (ICCV) 618–626 (2017). doi:10.1109/ICCV.2017.74.

35. Mathworks. Grad-CAM Reveals the Why Behind Deep Learning Decisions. MATLAB & Simulink https://www.mathworks.com/help/deeplearning/ug/gradcam-explains-why.html.

36. Sundararajan, M., Taly, A. & Yan, Q. Axiomatic Attribution for Deep Networks. http://arxiv.org/abs/1703.01365 (2017).

37. Erion, G., Janizek, J. D., Sturmfels, P., Lundberg, S. & Lee, S.-I. Improving performance of deep learning models with axiomatic attribution priors and expected gradients. http://arxiv.org/abs/1906.10670 (2020).

38. Amunts, K., Mohlberg, H., Bludau, S. & Zilles, K. Julich-Brain: A 3D probabilistic atlas of the human brain’s cytoarchitecture. Science (1979). 369, 988–992 (2020).

39. Dickscheid, T., et al. siibra-python. Preprint at 10.5281/zenodo.15591482 (2025).

40. Thomas Yeo, B. T., et al. The organization of the human cerebral cortex estimated by intrinsic functional connectivity. J. Neurophysiol. 106, 1125–1165 (2011).

41. Kahali, S., Raichle, M. E. & Yablonskiy, D. A. The Role of the Human Brain Neuron–Glia– Synapse Composition in Forming Resting-State Functional Connectivity Networks. Brain Sci. 11, (2021).

42. Baggio, H.-C. et al. Functional brain networks and cognitive deficits in Parkinson’s disease. Hum. Brain Mapp. 35, 4620–4634 (2014).

43. de la Fuente-Fernández, R. Imaging of dopamine in PD and implications for motor and neuropsychiatric manifestations of PD. Frontiers in Neurology vol. 4 JUL Preprint at 10.3389/fneur.2013.00090 (2013).

44. Droby, Amgad et al. Distinct Effects of Motor Training on Resting-State Functional Networks of the Brain in Parkinson’s Disease. Neurorehabil. Neural Repair 34, 795–803 (2020).

45. Pereira, J. B. et al. Initial cognitive decline is associated with cortical thinning in early Parkinson disease. Neurology 82, 2017–2025 (2014).

46. Rosenberg-Katz, K. et al. Gray matter atrophy distinguishes between Parkinson disease motor subtypes. Neurology 80, 1476–1484 (2013).

47. van den Berg, K. R. E., Johansson, M. E., Dirkx, M. F., Bloem, B. R. & Helmich, R. C. Changes in Action Tremor in Parkinson’s Disease over Time: Clinical and Neuroimaging Correlates. Movement Disorders 40, 292–304 (2025).

48. Taimur, M. et al. Frequency of Cognitive Impairment in Patients with Parkinson’s Disease. Cureus 11, e4733 (2019).

49. Inguanzo, A. et al. MRI subtypes in Parkinson’s disease across diverse populations and clustering approaches. NPJ Parkinsons Dis. 10, 159 (2024).

50. Zhang, F. et al. An anatomically curated fiber clustering white matter atlas for consistent white matter tract parcellation across the lifespan. Neuroimage 179, 429–447 (2018).

51. Pietracupa, S. et al. Understanding the role of cerebellum in early Parkinson’s disease: a structural and functional MRI study. NPJ Parkinsons Dis. 10, 119 (2024).

52. Gu, W., Bai, S. & Kong, L. A review on 2D instance segmentation based on deep neural networks. Image Vis. Comput. 120, 104401 (2022).

53. Ragguett, R. M., Eagleson, R. & de Ribaupierre, S. Evaluating normalized registration and preprocessing methodologies for the analysis of brain MRI in pediatric patients with shunt-treated hydrocephalus. Front. Neurosci. 18, (2024).

54. Despotović, I., Goossens, B. & Philips, W. MRI Segmentation of the Human Brain: Challenges, Methods, and Applications. Comput. Math. Methods Med. 2015, 450341 (2015).

